# UBE3A and transsynaptic complex NRXN1-CBLN1-GluD1 in a hypothalamic VMHvl-arcuate feedback circuit regulates aggression

**DOI:** 10.1101/2023.02.28.530462

**Authors:** Yi Nong, David C. Stoppel, Mark A. Johnson, Morgane Boillot, Jelena Todorovic, Jason Shen, Xinyu Zhou, Monica J.S. Nadler, Carrie Rodriguez, Yuda Huo, Ikue Nagakura, Ekkehard M. Kasper, Matthew P. Anderson

## Abstract

The circuit origins of aggression in autism spectrum disorder remain undefined. Here we report *Tac1*-expressing glutamatergic neurons in ventrolateral division of ventromedial hypothalamus (VMHvl) drive intermale aggression. Aggression is increased due to increases of *Ube3a* gene dosage in the VMHvl neurons when modeling autism due to maternal 15q11-13 triplication. Targeted deletion of increased *Ube3a* copies in VMHvl reverses the elevated aggression adult mice. VMHvl neurons form excitatory synapses onto hypothalamic arcuate nucleus AgRP/NPY neurons through a NRXN1-CBLN1-GluD1 transsynaptic complex and UBE3A impairs this synapse by decreasing *Cbln1* gene expression. Exciting AgRP/NPY arcuate neurons leads to feedback inhibition of VMHvl neurons and inhibits aggression. Asymptomatic increases of UBE3A synergize with a heterozygous deficiency of presynaptic *Nrxn1* or postsynaptic *Grid1* (both ASD genes) to increase aggression. Targeted deletions of *Grid1* in arcuate AgRP neurons impairs the VMHvl to AgRP/NPY neuron excitatory synapses while increasing aggression. Chemogenetic/optogenetic activation of arcuate AgRP/NPY neurons inhibits VMHvl neurons and represses aggression. These data reveal that multiple autism genes converge to regulate the VMHvl-arcuate AgRP/NPY glutamatergic synapse. The hypothalamic circuitry implicated by these data suggest impaired excitation of AgRP/NPY feedback inhibitory neurons may explain the increased aggression behavior found in genetic forms of autism.

**One Sentence Summary:** A feedback circuit in the hypothalamus that inhibits aggression is impaired by converging autism genetic defects.

## Introduction

### Autism spectrum disorders

(ASDs) are early onset behavioral disorders defined by impaired social communication, increased repetitive behaviors and restrictive interests. The condition is associated with variable comorbidities such as heightened irritability including aggressive behaviors. Heightened irritability can include excessive tantrums and self-injurious aggressive behaviors that often require medical treatment. One effective medication is risperidone, an agent that acts as an atypical antipsychotic providing one of the few FDA approved treatments for irritability behavior in ASD (*1*). However, the brain circuit mechanisms underlying elevated aggressive behaviors where medications might effectively modulate irritability in ASD remain undefined. Side effects of risperidone, obesity and dysregulated anterior pituitary function, correlate to major changes in gene expression that include up-regulated peptide hormone neurotransmitters such as NPY in the arcuate nucleus of the hypothalamus (*2*). Activity of neurons in the adjacent ventral lateral subdivision of ventral medial hypothalamus (VMHvl) is necessary and sufficient to promote attack behaviour in mice (*3–5*). The glutamatergic neurons of VMHvl include subsets that express estrogen receptor alpha (*Esr1*), progesterone receptor (*Pgr*), and neuropeptide substance P (*Tac1*). VMHvl *Esr1* and *Pgr* neurons were shown to promote aggressive behavior while a role for VMHvl *Tac1* neurons in regulating aggression is so far only suggested by a recent single cell-seq study showing increased expression of immediate early genes in response to aggressive behavior (*6*).

## Results

To study the role of VMHvl *Tac1* neurons in aggression, we applied the head mounted mini-microscope (Inscopix) technique to measure *in vivo* calcium dynamics in individual VMHvl *Tac1* neurons in freely moving mice (*7*). Calcium-sensitive fluorescent protein GCaMP7 was expressed selectively in VMHvl *Tac1* neurons by unilateral stereotactic injection of AAV9-hSyn-DIO-GCaMP7f virus into VMHvl of *Tac1-Cre* male mice. A gradient index lens was implanted in VMHvl after GCaMP7 virus injection (Fig. 1A and fig. S1, A and B). Recordings during the resident-intruder aggression behavior test revealed an increase of calcium events in VMHvl *Tac1* neurons during the introduction and attack of the intruder (Fig. 1A).

**Figure 1.**
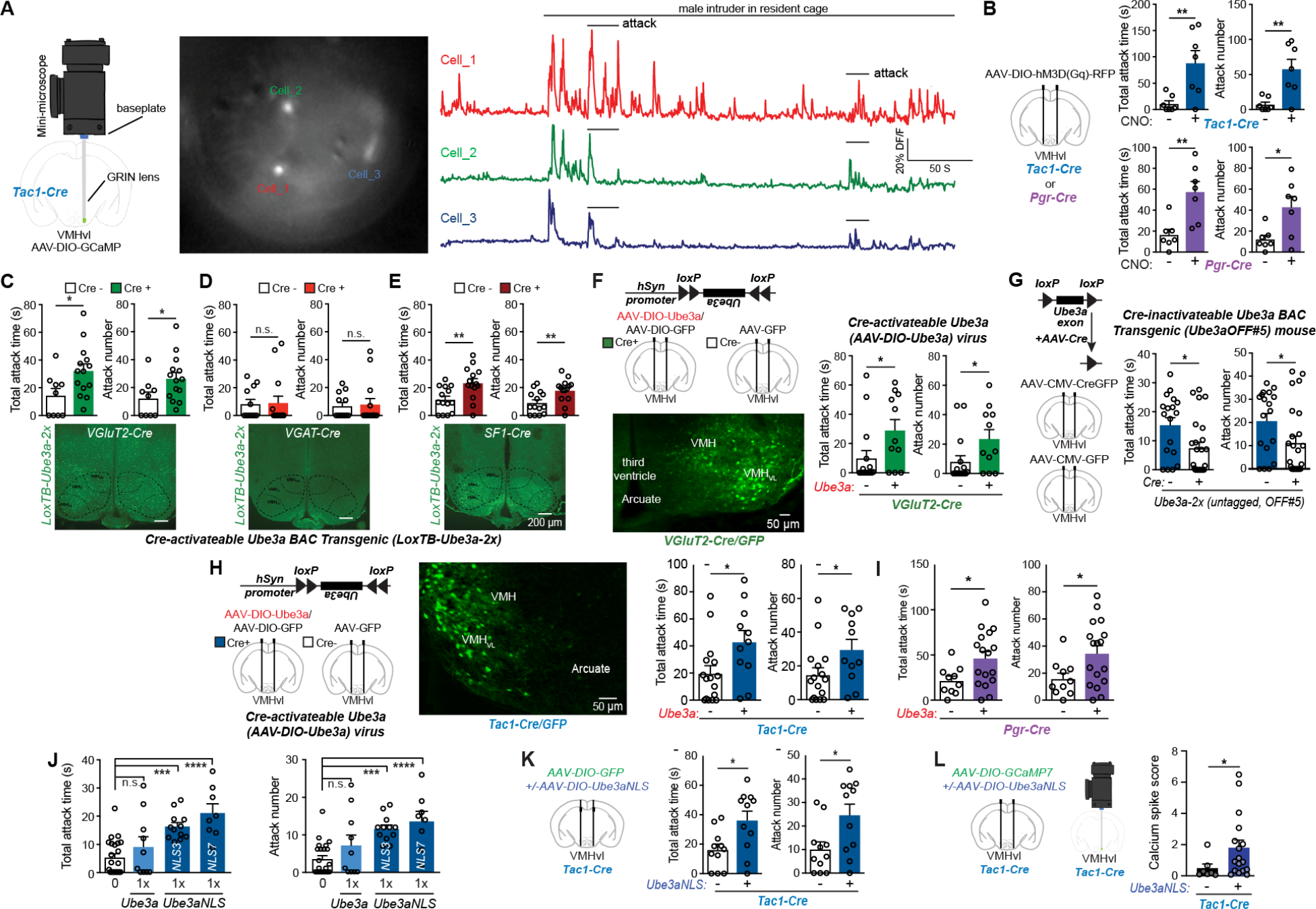
VMHvl *Tac1* neurons drive attack behavior and are the site where increased *Ube3a* gene dosage modeling a genetic ASD heightens aggression. (A) Representative images and traces of calcium events in three VMHvl *Tac1* neurons expressing GCaMP7 recorded using Inscopix miniature microscope with implanted GRIN lens when the resident male in its home-cage is exposed to a male intruder (resident-intruder paradigm). (B) Left: diagram of stereotactic injections of AAV virus in *Tac1-Cre* (*Tac1^tm1.1(cre)Hze^*) or *Pgr-Cre* (*Pgr^tm1.1(cre)Shah^*) mice. Right: upper panel: total attack time/number in *Tac1-Cre* mice (*n*=7) injected with AAV-DIO- hM3D(Gq)-RFP, comparing application of saline and CNO (1 mg/kg i.p.). *P* value of total attack time (*P_T_*) = 0.0068, and *P* value of attack number (*P_N_*) = 0.0042. Lower panel: total attack time/number in *Pgr-Cre* mice (*n*=7) injected with AAV-DIO-hM3D(Gq)-RFP, comparing application of saline and CNO (1 mg/kg i.p., *P_T_* = 0.0038, *P_N_* = 0.0165). (**C-E**) Total attack time and attack number in **C,** *VGluT2-Cre* (*Slc17a6^tm2(cre)Lowl^*):*LoxTB-Ube3a-2x* mice (*n* = 14) comparing *LoxTB-Ube3a-2x* littermates (*n* = 9) (*P_T_* = 0.026, *P_N_* = 0.042); **D**, *Vgat*-Cre (*Slc32a1^tm2(cre)Lowl^*)*:LoxTB-Ube3a-2x* mice (*n* =14) and *LoxTB-Ube3a-2x* littermates (*n* = 12) (*P_T_*= 0.9007, *P_N_* = 0.8203); **E**, *Sf1-Cre* (Tg(Nr5a1-cre)7Lowl):*LoxTB-Ube3a-2x* mice (*n* = 14) and *LoxTB-Ube3a-2x* littermates (*n* = 14) (*PT*= 0.0056, *P_N_* = 0.0085). Below, anti-FLAG antibody immunofluorescence in ventromedial hypothalamus (VMH), scale bars 200 μm. (**F**) Left: upper panel: diagram of construct of AAV-hSyn-DIO-Ube3a and stereotactic injections. Lower panel: representative image of anti-GFP antibody immunofluorescence in *VGluT2*-Cre mice injected with AAV-hSyn-DIO-Ube3a and AAV-hSyn-DIO-GFP in VMHvl, scale bar, 50 μm. Right: total attack time/number in *VGluT2*-Cre mice injected with AAV-hSyn-DIO- Ube3a:AAV-DIO-GFP (*n* = 16), compared to wild type littermates with *AAV-CMV-GFP* (*n* = 11, *P_T_* = 0.0373, *P_N_* = 0.0394) in VMHvl. (**G**) Left: diagram of construct of Cre-inactivate-able *Ube3a* BAC Transgenic (*Ube3a*OFF#5; figS2I, a conditional *Ube3a* transgene where *LoxP* site flank exons of the full-length untagged *Ube3a* gene) and stereotactic injections. Right: total attack time/number in *Ube3a*OFF#5 mice injected with AAV-CMV-CreGFP (*n* = 25) in VMHvl compared to AAV-CMV-GFP (*n* = 18, *P_T_* = 0.0146, *P_N_* = 0.0385). (**H**) Left: diagram of construct of AAV-hSyn-DIO-Ube3a, stereotactic injections. Middle: representative image of anti-GFP antibody immunofluorescence staining in *Tac1^Cre^* mice injected with AAV-hSyn-DIO- Ube3a and AAV-hSyn-DIO-GFP in VMHvl (scale bar, 50 μm). Right: total attack time/number in *Tac1-Cre* mice injected with AAV-hSyn-DIO-Ube3a:AAV-DIO-GFP (*n* = 11) in VMHvl, compared to wild type littermates injected with AAV-CMV-GFP (*n* = 16, *P_T_* = 0.025, *P_N_* = 0.0443). (**I**) Total attack time/number in *Pgr-Cre* mice injected with AAV-hSyn-DIO- Ube3a:AAV-DIO-GFP (*n* = 17) in VMHvl, compared to wild type littermates injected with AAV-CMV-GFP (*n* = 10, *P_T_* = 0.026, *P_N_* = 0.032). (**J**) Total attack time/number in wild type (*n* = 23) mice, comparing with *Ube3a-1x* (*n* = 10, *P_T_* = ns, *P_N_* =ns); with *Ube3a-NLS3-1x* mice (*n* = 12, *P_T_* < 0.0001, *P_N_* < 0.0001); and with *Ube3a-NLS7-1x* mice (*n* = 8, *P_T_* < 0.0001, *P_N_* < 0.0001). (**K**) Total attack time/number in *Tac1-Cre* mice injected with *AAV-DIO-GFP* + *AAV-DIO- Ube3a-NLS* (*n* =11) in VMHvl, compared to *AAV-DIO-GFP* (*n* = 11) (*P_T_*=0.0130, *P_N_*=0.0183). (**L**) Calcium events in VMHvl *Tac1* neurons recorded using Inscopix miniature microscope, compared *Tac1-Cre* mice injected with AAV9-hSyn-DIO-GCaMP7f (*n* = 7 cells) to AAV-DIO- Ube3a-NLS:AAV9-hSyn-DIO-GCaMP7f (*n* = 17 cells, *P*=0.0261; unpaired T-test with Welch’s correction). Unpaired two-tailed Student’s t-test was used to determine statistical significance when comparing two groups. Comparisons across two groups before and after CNO treatment analyzed by paired two-tailed Student’s t-test. Multiple groups were analyzed using 1-way ANOVA followed by Bonferroni’s Multiple Comparison Correction. Mean ± SEM shown, ns *P* > 0.05, \**P* < 0.05, \*\**P* < 0.01.

To test if VMHvl *Tac1* neuron activity is sufficient to drive attack behavior, we expressed chemogenetic tools in VMHvl *Tac1* neurons and activated these neurons during resident intruder testing (*8*). AAV-hSyn-DIO-hM3D(Gq)-mCherry virus was injected bilaterally into VMHvl of *Tac1-Cre* male mice (fig. S1C) to express excitatory Gq-coupled designer receptor *hM3D* activated by clozapine-N-oxide (CNO). As a positive control, *Pgr-Cre* expressing neurons in VMHvl were also tested. Administering CNO (1 mg/kg i.p.) dramatically increased total attack time and attack numbers in both *Tac1-Cre* and *Pgr-Cre* male mice when compared to animals receiving saline (Fig. 1B). In wild type male mice, CNO had no effect on attack behavior (fig. S1D). Similarly, in *Tac1-Cre* male mice injected with virus expressing only GFP protein, CNO had no effect on attack behavior (fig. S1, E and F). These results indicate augmented activity of VMHvl *Tac1* neurons is sufficient to magnify aggression.

There are now many genes implicated in ASD (*9, 10*) with recent studies suggesting the importance of polygenic interactions (*11, 12*). One of the most common strongly penetrant forms of genetic ASD results from increased dosages of the imprinted gene *UBE3A* (*9, 13, 14*). Maternal 15q11-13 triplications (due to a maternally-derived extranumerary isodicentric chromome 15q segment, idic15) triple the dosage of *UBE3A* gene expressed in mature neurons where the paternal allele is silenced. We studied *Ube3a* transgenic mice with additional copies of mouse full-length *Ube3a* gene designed to model this strongly penetrant genetic form of ASD (*10-12, 15-18*): *Ube3a- 1x* mice (heterozygous for the genomic *Ube3a* transgene insert) modeling maternal 15q11-13 interstitial duplication and *Ube3a-2x* mice (homozygous for the genomic *Ube3a* transgene insert) modeling maternal extranumerary isodicentric chromosome 15 (idic15) (fig. S2, A and B). *UBE3A* encodes an E3 ubiquitin ligase and transcriptional co-regulator protein (*14, 19–22*). We previously showed that increasing the dosage of this non-imprinted full-length *Ube3a* gene in mice (*Ube3a- 2x* mice) impairs social interactions and vocalizations while *Ube3a-1x* mice display more limited deficits (*14*). Male *Ube3a-2x* mice were found to display hightened levels of aggression frequently producing severe tail bites and occasional ulcerative hindlimb wounds on their co-housed wild type littermates (fig. S2D). In the resident-intruder test, a common laboratory test for aggression, we observed increased attack of the intruder by *Ube3a-2x* male mice compared to their wild type male littermates (fig. S2E).

To determine the neuronal subtypes where increased UBE3A might elevate aggression we generated a conditional *Ube3a*-*3xFLAG* mouse line (*LoxTB*-*Ube3a* mouse) where the extra copies of full-length *Ube3a* are expressed only in the presence of *Cre* recombinase. By inserting a *LoxP*- flanked translational blocker cassette (*LoxTB*) into intron 1 of the mouse *Ube3a* gene with a C- terminal FLAG tag (*Ube3a-3xFLAG*), the *LoxTB* cassette disrupts expression of all *Ube3a* splice variants. When Cre recombinase is expressed in specific cell types (through cell-type specific promoters), it deletes the *LoxTB* cassette, permitting cell-type specific expression of the FLAG- tagged UBE3A (fig. S2F). Male mice with the *Ube3a* transgene selectively expressed in glutamatergic neurons (*LoxTB-Ube3a-2x*: *VGluT2-Cre* mice, Fig. 1C, lower panel), but not in GABAergic neurons (*LoxTB-Ube3a-2x*:*Vgat*^Cre^ mice, Fig. 1D, lower panel), displayed increased attack towards the intruder when compared to littermates lacking Cre (Fig. 1, C and D, upper panel). No increase of attack behavior was observed in mice with *Ube3a* transgene selectively expressed in serotoninergic neurons (fig. S2G). The results indicate increasing *Ube3a* gene dosage in glutamatergic neurons is sufficient to heighten aggression.

To evaluate whether increases of *Ube3a* gene dosage in VMH might underlie this elevated aggression, we crossed *LoxTB-Ube3a-2x* and *Sf1-Cre* (steroidogenic factor 1 [*Sf1,Nr5a1*]) mice since *Sf1-Cre* is expressed predominantly in VMH (*23*). *LoxTB-Ube3a-2x*:*Sf1-Cre* mice displayed increased attack behavior compared to control littermate mice (Fig. 1E). To permit a spatial as well as cell-type specific assessment of the sites where increased *Ube3a* heightens aggression, we generated an AAV viral vector that expresses *Ube3a* in a Cre-dependent manner (AAV-hSyn-DIO-Ube3a, Fig.1F and fig. S2H). Some investigators have been concerned that a C-terminal epitope tage might impair normal functions of UBE3A and so we performed a series of studies including the use of AAV-hSyn-DIO-Ube3a where not C-terminal tage is added. Stereotactic co-injections of AAV-hSyn-DIO-Ube3a and AAV-hSyn-DIO-GFP into VMHvl of *VGluT2-Cre* adult male mice recapitulated the heightened aggression observed in *Ube3a*-2x mice when compared to AAV-hSyn-DIO-GFP alone (Fig. 1F, with GFP expression in VMHvl). The results suggest increases of UBE3A in glutamatergic neurons of VMHvl elevate aggression in an ongoing manner not contingent on developmental effects.

To test if the increased UBE3A in VMHvl is necessary for the increased aggression and whether removing the elevated UBE3A in adult mice can rescue this behavioral phenotype, we generated mice in which *LoxP* sites flanked exons of the full-length untagged *Ube3a* transgene (*Ube3a* OFF#5 mouse, Fig. 1G, left and fig. S2I). Again, note that the *Ube3a* OFF#5 mouse carries a *Ube3a* transgene lacking an epitope tag and also increases aggression. Strikingly, injection of AAV-CMV-CreGFP in VMHvl rescued the heightened aggression behavior observed in these adult *Ube3a*-2x (untagged) mice (Fig. 1G). The results indicate increases of UBE3A in VMHvl neurons is both sufficient and necessary for the heighten aggression seen in this genetic ASD mouse model and suggest reversibility of aggression in adulthood.

To determine if the aggression-promoting *Tac1* neuron in VMHvl might underlie this UBE3A-augmented aggression, AAV-hSyn-DIO-Ube3a (untagged) was stereotactically injected into VMHvl of *Tac1-Cre* male mice (Fig. 1, H and I). Progesterone-expressing neurons in VMHvl were also examined using *Pgr-Cre* male mice. Increasing UBE3A in either VMHvl *Tac1* or *Pgr* neurons increases attack behavior (Fig. 1, H and I).

We previously established that UBE3A acts as a potent transcriptional regulator through its actions in the neuronal cell nucleus in brain *in vivo* and others have shown co-activation with nuclear hormone receptors and zinc-finger *Sp1* transcription factors *in vitro* (*13, 21*). The same gene regulatory effects were observed with a C-terminal tagged and an untagged version of *Ube3a* transgene and reciprocal changes in many of the same genes was observed in mice where the maternal copy of the *Ube3a* gene was deleted. To test whether increases of UBE3A heighten aggression through its actions in the nucleus we used our mice where the extra gene copies of *Ube3a* carry a C-terminal fused nuclear localization signal. A 3xFLAG tag follows two tandem copies of a nuclear localization signal (*NLS*) in exon 12 of the mouse *Ube3a* transgene (fig. S2C). Comparing mice of either nuclear-targeted or non-targeted *Ube3a* transgene with equivalent mRNA and protein levels (*13, 14*), we found that mice with non-nuclear-targeted *Ube3a*-1x lacked the elevated aggression whereas two different founder *Ube3a-NLS*-1x mouse lines (“*Ube3a-NLS-*3 and / *Ube3a-NLS-*7” mice) displayed heightened aggression (Fig. 1J).

Therefore, the phenotype of nuclear-targeted *Ube3a-NLS*-1x mice is equivalent to the heightened aggression behavior observed in *Ube3a*-2x mice (non-nuclear-targeted). Targeting UBE3A to the nucleus selectively in VMHvl *Tac1*+ neurons, we observed that *Tac1-Cre* mice injected in VMHvl with AAV-DIO-Ube3a-NLS:AAV-DIO-GFP exhibited heightened aggression compared to littermate *Tac1-Cre* mice injected only with AAV-DIO-GFP alone (Fig. 1K and figs. S3, A and B). These mice also displayed pathologic aggression as the male mice expressing nuclear-targeted UBE3A not only attacked a male intruder, but also inflicted bite injuries on wild type female mice placed in their cages to prime them for aggression testing (fig. S3C). In contrast, when *Sf1-Cre* mice were injected with AAV-DIO-Ube3a-NLS:AAV-DIO-GFP in dorsal medial subdivision of ventromedial hypothalamus (VMHdm), no increase of aggression relative to littermate mice injected with AAV-DIO-GFP was observed (fig. S3, D to F), further supporting the role of VMHvl in aggression.

To test if nuclear-targeted UBE3A alters activity of VMHvl *Tac1* neurons during aggression behavior, we injected AAV9-hSyn-DIO-GCaMP7f into VMHvl of *Tac1-Cre* male mice with or without *AAV-DIO-Ube3a-NLS*. VMHvl *Tac1* neurons expressing Ube3a-NLS and AAV-DIO-GCaMP7f displayed increased calcium spiking activity when exposed to male intruder when compared to mice with AAV-DIO-GCaMP7f alone (Fig. 1L).

Nuclear-targeting of UBE3A magnifies its effects on gene regulation and we recently reported *Cbln1* is one of the genes most strongly repressed in cerebral cortex of *Ube3a*-2x and *Ube3a-NLS*-1x mice by quantitative RT-PCR (*13*). CBLN1 protein is a glutamate synapse organizer (*24*) and deleting *Cbln1* in ventral tegmental area (VTA) reduces synaptic transmission from VTA glutamatergic neurons to medium spiny neurons of nucleus accumbens while impairing sociability (*13*). *Cbln1* is also strongly expressed in VMH neurons where increased *Ube3a* amplifies aggression (Allen Brain Atlas: Mouse Brain). We found that *Cbln1* mRNA is repressed in VMH of *Ube3a-NLS7*-1x transgenic mice compared to wild type littermates (Fig. 2A). To further study how UBE3A regulates *Cbln1* gene expression, a 1.3 kb 5’ *Cbln1* promoter sequence was introduced to drive luciferase expression. When this *Cbln1* promoter-luciferase reporter was co-transfected with *Ube3a*-expressing or control vector, luciferase activity was reduced by as much as 54% by UBE3A (Fig. 2B). The results indicate UBE3A can repress the *Cbln1* promoter.

**Fig. 2.**
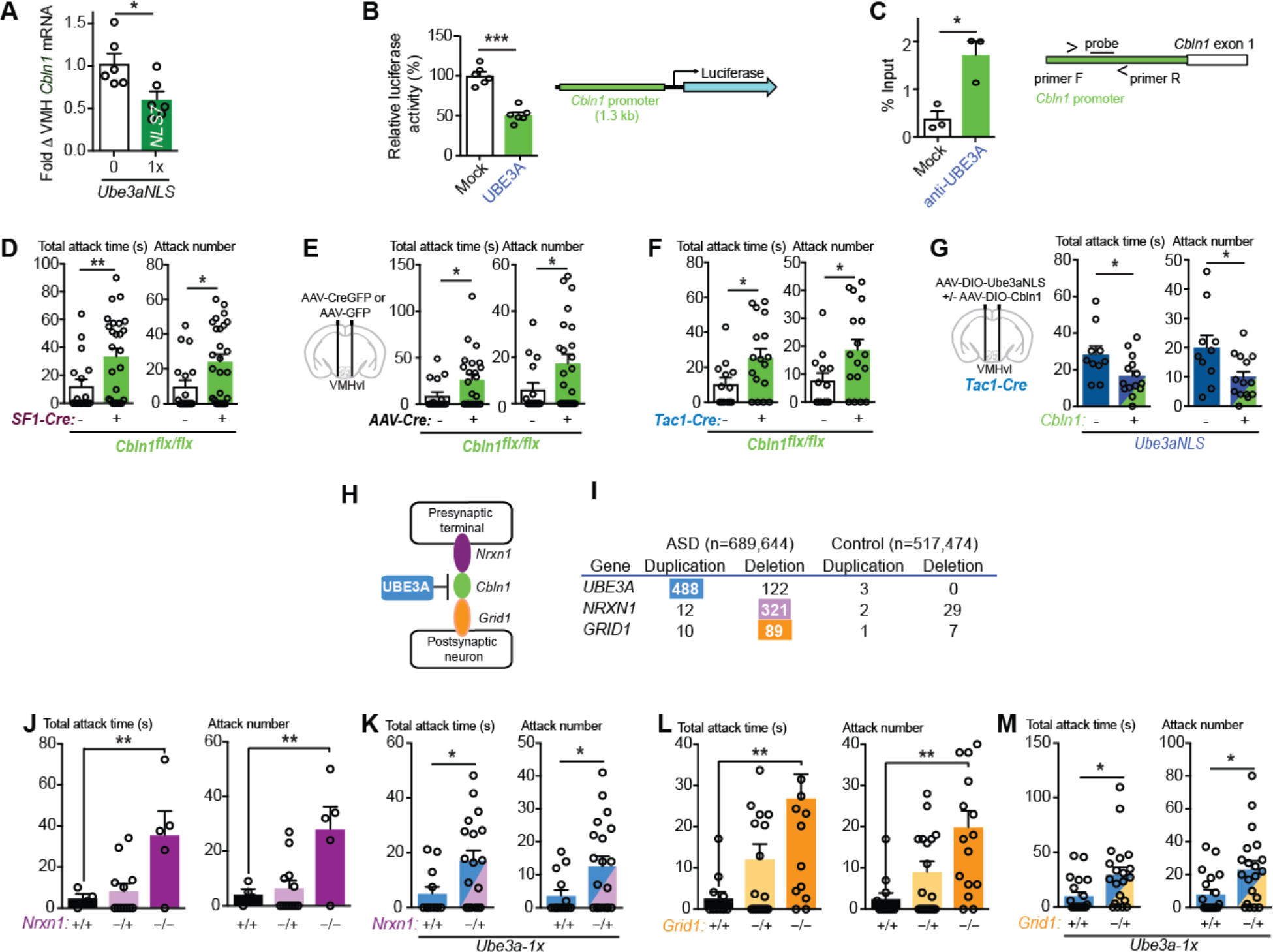
UBE3A represses *Cbln1* gene expression and synergizes with a loss of NRXN1- CBLN1-GRID1 transsynaptic complex members to promote aggression. (A) Quantitative RT-PCR of *Cbln1* mRNA in VMH in wild type littermate (*n* = 6), and *Ube3a-NLS-1x* transgenic mice (*n* = 6, *P* = 0.0317). (B) *Cbln1* 5’ promoter sequence (1.3 kb) driving luciferase reporter with co-transfected *Ube3a* (*n* = 6), comparing to mock plasmid (*n* = 6, *P* < 0.0001). Diagram shows the luciferase reporter construct. (C) The ratio of *Cbln1* promoter DNA immunoprecipitated from cortex samples, relative to input chromatin, compared anti-UBE3A antibody to “Mock” rabbit IgG (*n* = 3, *P* = 0.015). Diagram shows FAM/ZEN probe location. (**D**) Total attack time/number in mice with *Sf1*-*Cre*:*Cbln1^flx/flx^* (*n* = 26) compared to *Cbln1^flx/flx^* littermates (*n* = 18, *P_T_* = 0.0098, *P_N_* = 0.0180). (**E**) Total attack time/number in *Cbln1^flx/flx^* mice with injected AAV-CMV-CreGFP (*n* = 23) vs. AAV-CMV-GFP (*n* =16) in VMHvl (*P_T_* = 0.0418, *P_N_* = 0.0413). (**F**) Total attack time/number in mice with *Tac1-Cre*/*Cbln1^flx/flx^* (*n* = 17) compared to *Cbln1^flx/flx^* littermates (*n* = 13, *P_T_* = 0.0285, *P_N_* = 0.0313). (**G**) Total attack time/number in *Tac1-Cre* mice injected with *AAV-DIO-Ube3a-NLS* (*n* = 10), comparing with *AAV-DIO-Cbln1* + *AAV-DIO-Ube3aNLS* (*n* = 14, *P_T_* = 0.030, *P_N_* = 0.0245) in VMHvl. (**H**) Diagram of protein physical interactions between CBLN1, NRXN1, and GRID1. (**I**) Number of ASD cases with genomic copy number variations (CNVs) encompassing genes *UBE3A, NRXN1* and *GRID1* including duplications/triplications (Dup) and deletions (Del). (**J**) Total attack time/number in wild type littermates (n=4), *Nrxn1* heterozygous (*n* = 12, *P_T_* = ns, *P_N_* = ns) and *Nrxn1* homozygous deletion mice (*n* = 5, *P_T_* = 0.009, *P_N_* = 0.006). (**K**) Total attack time/number in mice with *Ube3a-1x* (*n* = 14), compared to *Ube3a-1x* and *Nrxn1* +/- (n = 20, *P_T_* = 0.0261, *P_N_* = 0.0257). (**L**) Total attack time/number in wild type littermates (*n* = 12), *Grid1* heterozygous (*n* = 22) and *Grid1* homozygous deletion mice (*n* = 16, *P_T_* = 0.0023, *P_N_* = 0.0018). (**M**) Total attack time/number in mice with *Ube3a-1x* (*n* = 20), compared to *Ube3a-1x* and *Grid1* +/- (*n* = 20, *P_T_* = 0.012, *P_N_* = 0.0085). An unpaired two-tailed Student’s t-test was used to determine statistical significance when comparing two groups. Mean ± SEM shown, ns *P* > 0.05, \**P* < 0.05, \*\**P* < 0.01, \*\*\**P* < 0.001.

We next performed chromatin immunoprecipitation (ChIP) from wild type mouse cortex using an anti-UBE3A antibody, and quantified the levels of UBE3A-bound *Cbln1* promoter sequences using quantitative PCR. The quantity of UBE3A-bound *Cbln1* promoter DNA relative to input chromatin is strongly enriched (1.72 ± 0.29) when compared to mock antibody (0.39 ± 0.16, Fig. 2C). The results suggest UBE3A functions as a transcriptional co-regulator physically interacting with and repressing the 5’ *Cbln1* promoter.

To determine if a loss of *Cbln1* in VMH is sufficient to increase aggression behavior, we combined homozygous floxed *Cbln1* (*Cbln1^flx/flx^*) mouse with *Sf1-Cre* to delete *Cbln1* in VMH neurons (fig. S3G). Male *Sf1-Cre*:*Cbln1^flx/flx^* mice displayed increased attack behavior when compared to *Cbln1^flx/flx^* littermates (Fig. 2D). Injecting AAV-CMV-CreGFP (vs. AAV-CMV-GFP) in VMHvl of adult *Cbln1^flx/flx^*mice also increased attack behavior (Fig. 2E). Furthermore, deleting *Cbln1* in *Tac1* neurons (male mice with *Tac1-Cre /Cbln1^flx/flx^* vs. *Cbln1^flx/flx^*) increased attack behavior (Fig. 2F). The results indicate the same brain regions and cell-types underlying the increased aggression evoked by increased UBE3A, which represses *Cbln1*, are also the site where *Cbln1* deletion increases aggression. To determine whether CBLN1 is the major target whereby UBE3A promotes aggression, we co-injected AAV-DIO-Ube3a-NLS in *Tac1-Cre* mice with or without AAV-hSyn-DIO-Cbln1. We found the increased aggression produced by nuclear-targeted UBE3A is reversed by an increase of CBLN1 in VMHvl *Tac1* neurons (Fig. 2G). These results confirm that UBE3A increases aggression through its actions in the nucleus of VMHvl *Tac1* neurons primarily by repressing *Cbln1* gene expression.

We previously used a bioinformatics strategy to examine the physical interactions between the proteins encoded by UBE3A-regulated and other ASD genes and discovered that CBLN1 physically interacts with two other gene products frequently deleted in ASD, presynaptic NRXN1 (encoded by *NRXN1* gene,) and postsynaptic GluD1 (encoded by gene *GRID1,* Fig. 2H). CBLN1 organizes glutamate synapses by binding to NRXN1 and GluD1. This NRXN1-CBLN1-GluD1 transsynaptic trimolecular complex regulates glutamate synapse formation and maintenance (*24*). Amongst *NRXN* and *GRID* gene family members, heterozygous deletions encompassing *NRXN1* were found in 321 cases (second most frequent CNV after *UBE3A* in some case series) and *GRID1* in 89 cases (Fig. 2I, AutDB (*25*)). *UBE3A* gene copy numbers are likely underestimated as autism cases diagnosed with extranumerary isodicentric chromosome (idic15) by cytogenetics are often excluded.

We found that homozygous deletion of CBLN1 binding partner genes *Nrxn1* or *Grid1* heightens aggression in mice as previously reported (*26, 27*), whereas heterozygous *Nrxn1* or *Grid1* deletions does not (Fig. 2, J to M), similar to the behavioral effects of *Ube3a*-1x (Fig. 1J). Since *Ube3a*-1x mice does moderately repress *Cbln1* expression (*13*), we reasoned that haploinsufficiency of *Grid1* or *Nrxn1* might mimic the effects of homozygous deletion by enhancing aggression when added to the partial *Cbln1*-repressed state of the *Ube3a*-1x mice. Remarkably, heterozygous deletion of *Nrxn1* or *Grid1* increased aggression in an *Ube3a*-1x “genetic background” (Figs. 2, K and M) providing a biological *proof-of-concept* example whereby ASD-linked genetic defects could display polygenic interactions to effect behavior (*10-12, 15-18*).

The decreased expression of glutamatergic synapse organizer *Cbln1* that results from increased UBE3A suggested *Ube3a-2x* mice might display impaired activation of a neuron population targeted by VMHvl neuron glutamate synapses during aggression behavior. To study the neuronal activity changes during aggression behavior in *Ube3a-2x* and littermate mice, we counted c-fos positive neurons in select brain regions immediately after aggression behavior testing (fig. S4A). We focused on brain regions which synaptically connect to VMHvl according to a previous report (*28*) and found changes in c-fos in several of these brain regions in *Ube3a-2x* mice (Fig. 3, A to F and fig. S4, B to G). The number of c-fos positive neurons was increased in brainstem outflow pathways of VMHvl including periaqueductal grey (PAG, mostly in the caudal region, Figs. 3, A and C) and dorsal raphe (fig. S4, D and E). Increased numbers of c-fos positive neurons were also observed in a specific medial ventral subregion of VMHvl (Fig. 3, B and F).

**Fig. 3.**
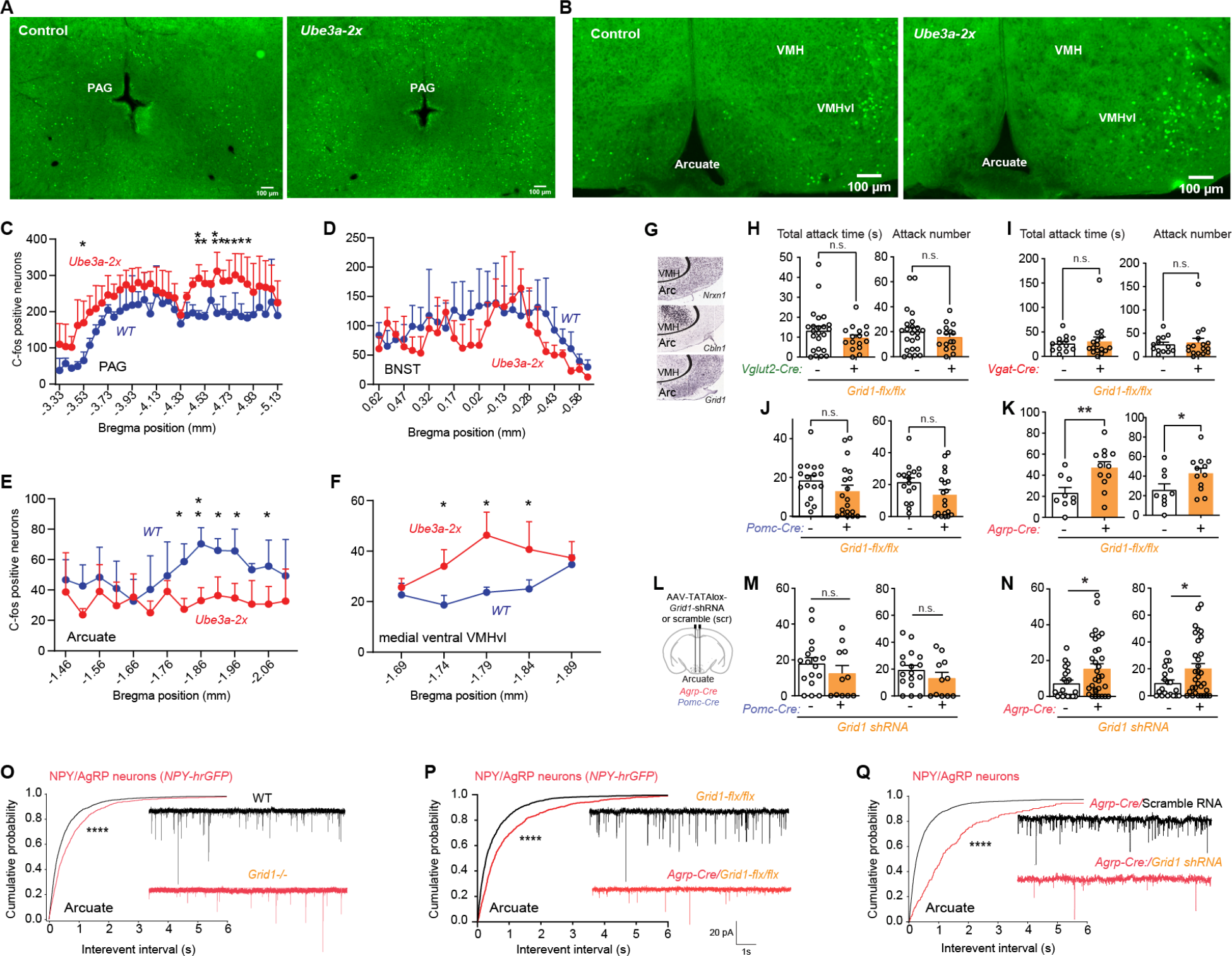
Loss of postsynaptic glutamate receptor delta subunit gene *Grid1* in arcuate AgRP/NPY neurons impairs their glutamatergic synapses to increase aggression. (A) Representative image of anti-c-fos antibody immunofluorescence staining in PAG after aggression behavior test compared *Ube3a-2x* mice with their little mate control (scale bar, 100 μm). (B) Representative image of anti-c-fos staining in arcuate and VMHvl compared *Ube3a-2x* mice with control (scale bar, 100 μm). C, D, E, F, Quantitatively counting c-fos positive neurons in PAG (C, *P* = 0.0396); BNST (D, *P* = 0.4578); arcuate (E, *P* = 0.0249) and a selection of region of VMHvl (**F**, *P* = 0.0472) after aggression behavior test compared *Ube3a-2x* mice with their littermate control. (**G**) *In situ* hybridization of parasagittal VMH-arcuate hypothalamus sections probed for *Nrxn1*, *Cbln1*, and *Grid1* mRNA (adapted from Allen Mouse Brain Atlas (2004)). (**H**) Total attack time/number in mice with *Vglut2-Cre*:*Grid1^flx/flx^* (*n* = 16) compared to *Grid1^flx/flx^* littermates (*n* = 23, *P_T_* = 0.2612, *P_N_* = 0.3579). (**I**) Total attack time/number in mice with *Vgat-Cre*:*Grid1^flx/flx^* (*n* = 13) compared to *Grid1^flx/flx^* littermates (*n* = 16, *P_T_* = 0.7061, *P_N_* = 0.7208). (**J**) Total attack time/number in mice with *Pomc-Cre* (Tg(Pomc1-cre)16Lowl):*Grid1^flx/flx^* (*n* = 19) compared to *Grid1^flx/flx^*littermates (*n* = 17, *P_T_* = 0.1682, *P_N_* = 0.0753). (**K**) Total attack time/number in mice with *Agrp-Cre* (*Agrp^tm1(cre)Lowl^*):*Grid1^flx/flx^* (*n* = 12) compared to *Grid1^flx/flx^* littermates (*n* = 9, *P_T_* = 0.0067, *P_N_* = 0.0498). (**L**) Diagram of stereotactic injections. (**M**) Total attack time/number in *Pomc-Cre* mice injected with AAV-TATAlox-Grid1-shRNA (*n* = 11) in arcuate, comparing to WT mice injected with AAV-TATAlox-Grid1-shRNA (*n* = 17, *P_T_* = 0.3212, *P_N_* = 0.272). (**N**) Total attack time/number in *Agrp-Cre* mice injected with AAV-TATAlox-Grid1-shRNA (*n* = 32) in arcuate, comparing to mice injected with AAV-TATAlox-scrambled-shRNA (*n* = 20, *P_T_* = 0.0403, *P_N_* = 0.0360). (**O**) Cumulative frequency plot of miniature excitatory post-synaptic current (mEPSC) inter-event intervals in arcuate NPY GFP^+^ neurons in mice with homozygous deletion *Grid1* (*n* = 4 mice, 16 cells) and control mice (*n* = 5 mice, 14 cells), Kolmogorov-Smirnov (KS) test, *P* < 0.00001. (**P**) Cumulative frequency plot of miniature excitatory post-synaptic current (mEPSC) inter-event intervals in arcuate NPY GFP^+^ neurons (labeled by *Npy-hrGFP* allele; B6.FVB-Tg(Npy-hrGFP)1Lowl/J) in mice with *Grid1* knockout (*Agrp-Cre*:*Grid1*^flx/flx^) in arcuate Agrp/NPY neurons (*n* = 6 mice, 11 neurons) compared to control wild type mice (*n* = 4 mice, 14 cells,). (**Q**) Cumulative frequency plot of miniature excitatory post-synaptic current (mEPSC) inter-event intervals in arcuate NPY GFP^+^ neurons (labeled by *Npy-hrGFP* allele) in mice with *Grid1* knockdown (AAV-hSyn-TATAlox-Grid1-shRNA vs. scrambled) in arcuate Agrp/NPY neurons (*n* = 3 mice, 6 neurons) compared to control wild type mice (*n* = 6 mice, 18 cells). An unpaired two-tailed Student’s t-test was used to determine statistical significance when comparing two groups. Multiple groups (Fig. **3C-F**) are analyzed using repeated two-way ANOVA with Multiple Comparison. mEPSC inter-event intervals and amplitude are analyzed using Kolmogorov-Smirnov (KS) test. Mean ± SEM shown, ns *P* > 0.05, \**P* < 0.05, \*\**P* < 0.01, \*\*\**P* < 0.001, \*\*\*\**P* < 0.0001. See also fig. S5 and S6.

There were no changes in c-fos expression observed in the bed nucleus of the stria terminalis (BNST) (Fig. 3D), ventral premamillary nucleus (PMv), or the full VMHvl region (fig. S4, B, C). Medial amygdala (MEA), by contrast, showed a decrease in the number of c-fos positive neurons in the caudal region (fig. S4, F and G). Interestingly, the number of c-fos positive neurons was also decreased in arcuate nucleus in *Ube3a-2x* mice (Fig. 3, B and E).

**Fig. 4.**
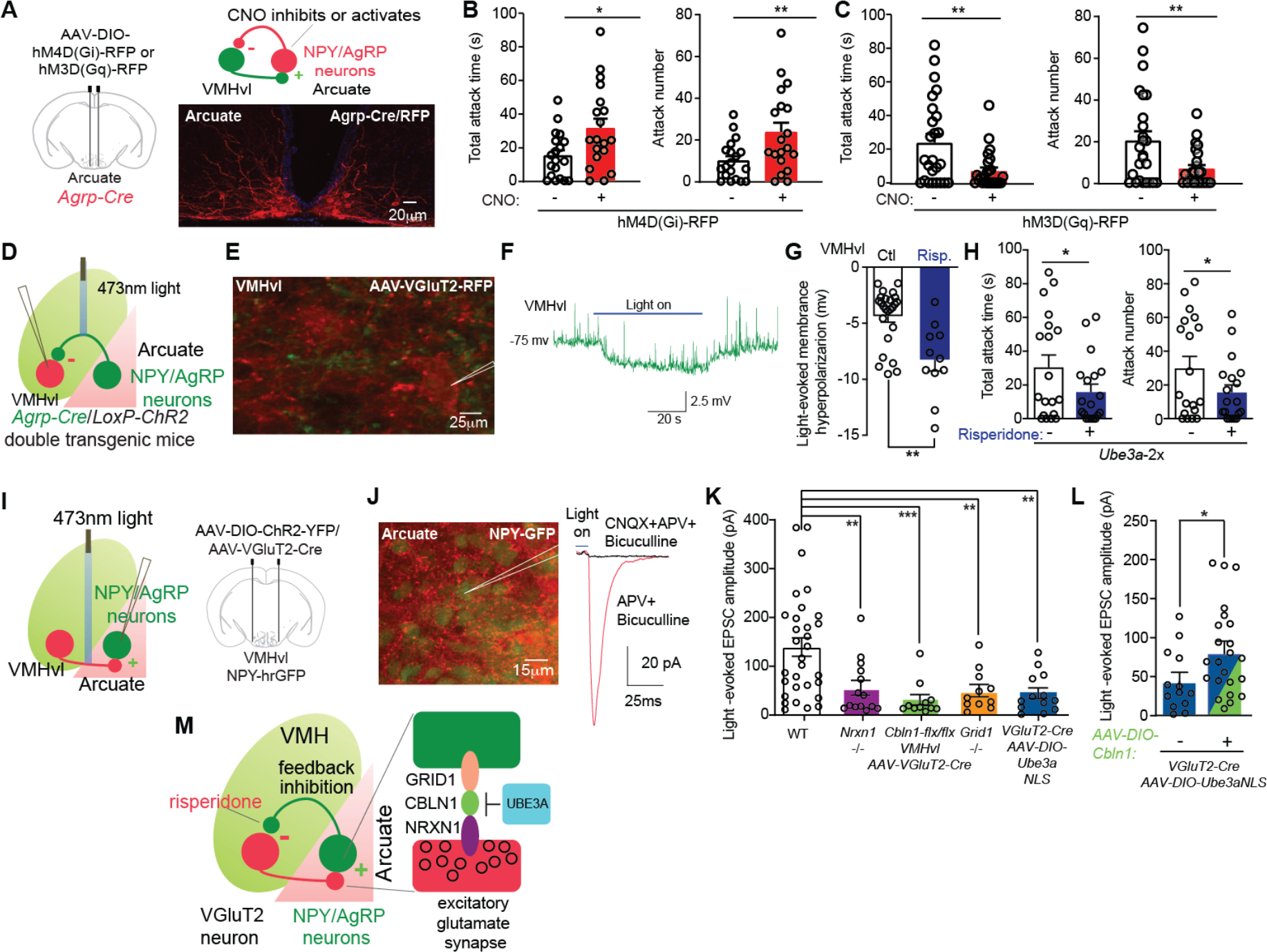
NRXN1-CBLN1-GRID1 transsynaptic complex forms excitatory synapses from VMHvl to arcuate NPY/AgRP neurons which provide feedback inhibition to VMHvl that is augmented by aggression medication risperidone. (A) Left: diagram of stereotactic injections. Right: representative image of anti-DsRed antibody immunofluorescence staining in *Agrp-Cre* mice injected with AAV-DIO-hM4D(Gq)-RFP in arcuate (scale bar, 30 μm). (B) Total attack time/number in *Agrp-Cre* mice with AAV-DIO-hM4D(Gi)-RFP injected in arcuate, comparing application of saline with CNO (1 mg/kg i.p., *n* = 19, *P_T_* = 0.0016, *P_N_* = 0.001). (C) Total attack time/number in *Agrp-Cre* mice with AAV-DIO-hM3D(Gq)-RFP injected in arcuate, comparing application of saline with CNO (1 mg/kg i.p., *n* = 25, *P_T_* = 0.0035, *P_N_* = 0.0062). (D) Diagram of VMHvl-arcuate circuit with light-excitation of arcuate AgRP neuron axon terminals expressing ChR2. (**E**) Representative image of AgRP/NPY axons (green, *Npy-hrGFP* transgenic mice) surrounding VMHvl neurons (red, labelled by AAV-VGlut2-mCherry, scale bar 15 μm). (**F**) Representative trace of recorded VMHvl neuron hyperpolarized by light-stimulated arcuate AgRP/NPY neuron axons. (**G**) Amplitude of light-evoked hyperpolarization in VMHvl neurons with stimulation of arcuate neurons axons in control wild type mice (*n* = 6 mice, 25 neurons, -4.40 ± 0.50 mV), or mice treated with risperidone (*n* = 4 mice, 11 neurons, -8.29 ± 0.98 mV). *P* = 0.0030 for control vs. risperidone treatment. (**H**) Total attack time/number in *Ube3a-2x* mice with risperidone treatment (1 mg/kg i.p., *n* = 25) comparing to saline treatment *(n* = 24, *P_t_* = 0.0475, *P_N_* = 0.0291). (**I**) Diagram of VMHvl-arcuate circuit with light-excitation of VMHvl glutamatergic neuron axon terminals. (**J**) Left: Diagram of stereotactic injections; and a representative image showing that labeled AgRP/NPY neurons (green, *Npy-hrGFP* mice) are surrounded by ChR2- expressing axons from VMHvl glutamate neurons (red, scale bar 15 μm). Right: representative trace of light-evoked AMPA excitatory post-synaptic currents (EPSCs) recorded in arcuate NPY GFP^+^ neurons by stimulating VMHvl terminals projecting to arcuate NPY neurons. (**K**) Amplitude of light-evoked EPSCs recorded in arcuate NPY GFP^+^ neurons when stimulating VMHvl ChR2- expressing terminals projecting to arcuate NPY neurons in mice with homozygous *Nrxn1* deletion (*n* = 5 mice, 14 neurons), *Grid1* deletion (*n* = 6 mice, 11 cells), *Cbln1* deletion in VMHvl (*(n* = 4 mice, 13 neurons) or increased nuclear UBE3A (VGluT2-Cre mice with AAV-hSyn-DIO-Ube3a- NLS in VMHvl) (*n* = 4 mice, 13 neurons); compared to control wild type mice (*n* = 8 mice, 31 cells). The *P* value of one-way ANOVA is <0.0001; the *P* values of Bonferroni’s multiple comparisons test are: *P* = 0.0039 for *Nrxn1* KO vs. WT; *P* = 0.0004 for *Cbln1* KO vs. WT; *P* = 0.0047 for *Grid1* KO vs. WT; *P* = 0.0012 for increasing nuclear *UBE3A* vs. WT. (**L**) Amplitude of light-evoked EPSCs recorded in arcuate NPY GFP^+^ neurons by stimulating VMHvl terminals projecting to arcuate NPY neurons in mice injected with AAV-hSyn-DIO-Ube3a-NLS in VMHvl (*n* = 4 mice, 13 neurons) compared to AAV-hSyn-DIO-Cbln1 + AAV-hSyn-DIO-Ube3aNLS (*n* = 7 mice, *n* = 23 neurons, *P* = 0.0409). (**M**) Model where VMHvl neurons form excitatory glutamatergic synapses onto arcuate NPY/AgRP neurons, which provide feedback inhibition by hyperpolarizing VMHvl neurons, serving as a gate to limit aggression. Paired two-tailed Student’s t-test was used for comparisons across two groups before and after CNO treatment in panel **B**, **C.** Multiple groups (panel **K**) are analyzed using one-way ANOVA with Bonferroni’s Multiple Comparison Correction. Unpaired two-tailed Student’s t-test was used to determine statistical significance of panel **G**, **H**, **I**. Mean ± SEM shown, ns *P* > 0.05, \**P* < 0.05, \*\**P* < 0.01, \*\*\**P* <0.001.

VMHvl neurons receive multiple inputs, and project in turn to multiple brain regions. The arcuate nucleus is one of six brain regions harboring the most input-output connections with VMHvl neurons (*28*). *Cbln1 mRNA* and *Nrxn1 mRNA* are highly expressed in VMHvl neurons, while, in hypothalamus, *Grid1 mRNA* is enriched in arcuate nucleus, in addition to its expression in multiple other brain regions (Allen Brain Atlas-mouse, Fig. 3G). These observations led us to reason that the increased aggression resulting from VMHvl-targeted loss of CLBN1 might disrupt NRXN1-CBLN1-GRID1 transsynaptic complexes to reduce activation of arcuate nucleus during aggression in *Ube3a-2x* mice. Consistent with this idea, a previous study found silencing neuropeptide F-expressing neurons (identified as the fly analog of NPY) increases aggression in fly (*29*), and chemogenetic activation of arcuate NPY/AgRP neurons decreases aggression in mice (*30*). Based on these observations, we inferred that a loss of *Grid1* specifically in arcuate nucleus might impair its excitatory synaptic inputs to decrease its activity leading to increased aggression. To test this hypothesis, we generated *Grid1^flx/flx^* mice and combined them with several mouse promoter-Cre lines. Deleting *Grid1* in glutamatergic (*Vglut2^Cre^*:*Grid1^flx/flx^*) or GABAergic (*Vgat-Cre*:*Grid1^flx/flx^*) neurons failed to increase aggression (Figs. 3, H and I). Similarly, deleting *Grid1* in POMC neurons (*Pomc-Cre*:*Grid1^flx/flx^*), a major neuron subtype in arcuate nucleus, failed to increase aggression (Fig. 3J). In contrast, deleting *Grid1* in AgRP neurons (*Agrp-Cre*:*Grid1^flx/flx^*) increased aggression (Fig. 3K). To test if reducing *Grid1* specifically in AgRP neurons of the hypothalamic arcuate nucleus increases aggression, we generated an AAV viral vector that expresses *Grid1-shRNA* in a Cre-dependent manner. Expressing *Grid1-shRNA* in arcuate AgRP neurons (*Agrp-Cre*) but not in arcuate POMC neurons (*Pomc-Cre*) increased aggression when injecting AAV-hSyn-TATAlox-Grid1-shRNA (vs. AAV-hSyn-TATAlox-scrambled-shRNA) (Fig. 3, L to N). Thus, a loss of CBLN1’s postsynaptic binding partner gene *Grid1* in arcuate AgRP/NPY neurons, but not in other major neuronal cell-types, recapitulates the aggression-promoting effects of germline *Grid1* deletion.

*Cbln1-*expressing glutamatergic cerebellar granule cell neurons form excitatory synapses onto local inhibitory neurons in a *Grid1*-dependent manner (*31*). Caged glutamate photostimulation of the VMHvl region elicits excitatory synaptic responses in arcuate NPY/AgRP neurons (*32*). Therefore, we tested whether a loss of *Grid1* in arcuate NPY/AgRP neurons impairs glutamate synaptic transmission onto NPY/AgRP neurons. Whole–cell patch clamp recordings of arcuate NPY/AgRP neurons were guided using the *Npy-hrGFP* transgene (fig. S5, A to C). Miniature excitatory postsynaptic currents (mEPSCs) were recorded in the presence of bath voltage-gated sodium channel blocker tetrodotoxin (TTX) and GABA_A_ receptor antagonist bicuculline. Loss of *Grid1* reduced mEPSC frequency in arcuate NPY/AgRP neurons in homozygous *Grid1* knockout mice with *Npy-hrGFP* (Fig. 3O) with no change in mEPSC amplitude (fig. S5D). Deletion of *Grid1* in AgRP neurons (*Agrp-Cre*:*Grid1^flx/flx^*) or knockdown of *Grid1* in arcuate AgRP neurons (AAV-hSyn-TATAlox-Grid1-shRNA vs. scrambled) also reduced mEPSC frequency (Fig. 3, P and Q) with no change in mEPSC amplitude (fig. S5, E). Thus, three genetic interventions that increase aggression behavior concurrently impair glutamate synapses onto arcuate NPY/AgRP neurons.

To directly examine whether decreasing arcuate AgRP/NPY neuron activity increases aggression, *Cre*-inducible inhibitory AAV-hSyn-DIO-hM4D(Gi) was injected into the arcuate nucleus of *Agrp-Cre* male mice (Fig. 4A). Applying the *hM4D* ligand CNO (vs. saline, i.p.) to inhibit these neurons increased attack behavior in these mice (Fig. 4B). In contrast, activating arcuate AgRP/NPY neurons decreased attack behavior using excitatory AAV-hSyn-DIO-hM3D (Gq) injected into the arcuate nucleus of *Agrp-Cre* male mice and applying the *hM3D* ligand CNO (vs. saline, i.p.) (Fig. 4C). Injecting AAV-hSyn-DIO-GFP into these mice and applying CNO failed to change aggression behaviors (fig. S5, G and H). The results indicate that arcuate NPY/AgRP neuron activity is necessary and sufficient to inhibit aggression.

Activity of the inhibitory arcuate NPY/AgRP neurons could inhibit target neurons such as those in VMHvl to decreased aggression. To study the effects of stimulating Agrp/NPY neuron axons on activity of VMHvl neurons, we applied the optogenetics technique to specifically excite NPY/AgRP neuron axon terminals expressing light-activated excitatory opsin in *Agrp-Cre* crossed to *Ai32* mice (*33*). Ai32 mice contain a *Cre-*inducible allele encoding the *Channelorhodopsin-2* (*ChR2*) (*34*). We found the membrane potential of VMHvl neurons was hyperpolarized by -4.40 ± 0.50 mV when ChR2-expressing NPY/AgRP neuron axon terminals were stimulated by light (Fig. 4, D to G). This result establishes that arcuate NPY/AgRP neurons inhibit VMHvl neurons.

Strikingly, chronic risperidone treatment (1 mg/kg twice daily i.p., for 15 consecutive days) magnified the NPY/AgRP axon terminal hyperpolarization of VMHvl neurons to -8.29 ± 0.98 mV (Fig. 4G). This same risperidone treatment regimen reduced the heightened aggression in *Ube3a*- 2x male mice (Fig. 4H). These results reveal direct evidence that arcuate NPY/AgRP axon terminals provide synaptic inhibition to VMHvl neurons, and suggest that augmenting such arcuate NPY/AgRP inhibition might be a novel therapeutic approach to alleviating increased aggressive behaviors.

Loss of *Grid1* impaired glutamatergic synaptic transmission onto arcuate NPY/AgRP neurons. We examine whether a loss of other components of the NRXN1-CBLN1-GRID1 transsynaptic complex or an increase of nuclear UBE3A also impairs such synapses. mEPSC frequency of arcuate NPY neurons was reduced in mice with homozygous *Nrxn1* deletion, *Cbln1* deletion in VMH *(Sf1-*Cre:*Cbln1^flx/flx^*), or increased nuclear UBE3A (*Ube3a-NLS7* mice) with no change in mEPSC amplitude (fig. S6, A to F).

The mEPSCs recorded in arcuate NPY/AgRP neurons could come from a variety of afferent glutamatergic neurons. To measure synaptic transmission selectively from VMHvl glutamate neurons to arcuate NPY/AgRP neurons we deployed optogenetics to selectively photo- excite ChR2-expressing VMHvl glutamate neuron axon terminals when AAV-hSyn-DIO-ChR2 with AAV-VGlut2-mCherry-2A-Cre were co-injected into VMHvl (Fig. 4I and fig. S6, G to H). Whole–cell patch-clamp recordings were performed in GFP-labeled arcuate NPY/AgRP neurons in *Npy-hrGFP* transgene mice 4 week after virus injection. The light-evoked excitatory post- synaptic currents (eEPSCs) recorded in the presence of antagonists of GABAA (bicuculline) and NMDA (D-2-amino-5-phosphonovaleric acid, D-APV) receptors were blocked by AMPA ionotropic glutamate receptor antagonist CNQX (6-cyano-7-nitroquinoxaline-2,3-dione, 30 μM, Fig. 4J). Compared to controls, the light-evoked AMPA EPSC amplitude was strongly reduced in mice with homozygous *Nrxn1* or *Grid1* deletion (Fig. 4K). Increasing nuclear UBE3A in VMHvl (*via* co-injection of AAV-Ube3a-NLS with VGluT2-Cre and AAV-hSyn-DIO-ChR2), or deleting CBLN in VMHvl (*via* co-injection of AAV-VGluT2-Cre and AAV-hSyn-DIO-ChR2 in VMHvl of *Cbln1^flx/flx^* mice) also markedly reduced light-evoked AMPA EPSC amplitude (Fig. 4K). Strikingly, adding CBLN1 (*via* co-injection of AAV-hSyn-DIO-Cbln1 with AAV-Ube3a-NLS) recovered the evoked EPSC amplitude repressed by the increased nuclear UBE3A (Fig. 4L). The results indicate glutamatergic synapses from VMHvl to arcuate NPY/AgRP neurons depend on each component of the NRXN1-CBLN1-GRID1 transsynaptic complex. Increased UBE3A promotes aggression by acting in the nucleus to impair this synapse largely by decreasing CBLN1.

## Discussion

Taken together, these results led us to propose a local circuit model where VMHvl glutamatergic neurons that express *Ube3a*, *Nrxn1* and *Cbln1* form excitatory glutamatergic synapses onto adjacent arcuate NPY/AgRP neurons that express *Grid1* (Fig. 4M). In this model, VMHvl neurons form glutamatergic synapses organized by NRXN1-CBLN1-GluD1 transsynaptic complex onto arcuate NPY/AgRP neurons which then project back to inhibit VMHvl neurons. This feedback loop acts as a gate to limit aggressive behavior. The findings reveal a distinct microcircuitry defect as a basis for elevated aggressive behavior across multiple genetic forms of ASD. The genetic defects exert their effects by disrupting glutamatergic synapses from VMHvl to arcuate NPY/AgRP which in turn are needed to activate these feedback inhibitory neurons (Fig. 4M).

Irritability/aggressive behaviors seen in autism are a frequent reason for pharmacological treatment with antipsychotics such as risperidone, yet these medications have a range of behavioral and metabolic side effects (*35, 36*). Here we show that risperidone represses the elevated aggression of *Ube3a-2x* mice while increasing the magnitude of arcuate AgRP/NPY mediated hyperpolarization of VMHvl neurons. This result suggests that by augmenting AgRP/NPY neuron synaptic transmission risperidone might both repress aggression while potentially also increasing feeding to cause weight gain (a known side effects). Other more complex pathways should also be considered. A study reported arcuate NPY/AgRP neurons may repress aggression through their projections to medial amygdala neurons with secondary projections to the bed nucleus of stria terminalis (*30*). Other neuronal pathways have also been identified to inhibit aggression behavior including the subparaventricular zone GABAergic neurons that transduce circadian signals (*37*) and the lateral septum GABAergic neurons (*38*).

### Future studies could examine if risperidone also augments arcuate NPY/AgRP neuron synaptic transmission to other targets such as the medial amygdala

We establish that an array of converging genetic defects including increases of UBE3A or a loss of NRXN1-CBLN1-GRID1 transsynaptic complex members that are impacted by ASD-associated copy number variations impair excitatory glutamatergic synapses from VMHvl to arcuate NPY/AgRP neurons while increasing aggressive behaviors. Heterozygous *NRXN1* or *GRID1* deficiencies as found in some cases of ASD increase aggression in mice only when combined with moderate increases of Ube3a dosage (*Ube3a-1x* mice), providing a biological proof-of-concept example for how genetic interactions could give rise to behavioral deficits in neuropsychiatric diseases by acting on multiple members of a molecular pathways such as a transsynaptic complex. Importantly, the elevated aggression caused by increases of UBE3A can be rescued by viral vector-based *Cbln1* expression in VMHvl neurons. Also removing the elevated UBE3A expression in VMHvl during adulthood rescues the heightened aggression.

Behavioral reversibility in adulthood highlights the plasticity of this circuits and implicates its potential as a future therapeutic target to treat irritability and aggressive behaviors in ASD. Our findings, that multiple genetic forms of ASD disrupt VMHvl-arcuate AgRP/NPY neuron glutamate synapses and that augmenting the aggression-limiting feedback inhibition from arcuate to VMHvl reduces aggression pave the way for precision medicine approaches specifically directed at this neuronal circuitry.

## Materials and Methods

### Animals

The Harvard Medical Area Standing Committee on Animals and the Institutional Animal Care and Use Committee of Beth Israel Deaconess Medical Center approved all mouse protocols. Mice were housed at the Center for Life Sciences barrier animal facility in sex-matched groups of 3-5 with ad libitum food and water access. Unless otherwise specified, littermate controls were used and cohorts were male only. Ube3a transgenic mice, *Ube3aNLS* mice and *LoxTB-Ube3a* mice were generated using bacterial artificial chromosome (BAC) recombineering techniques as described previously (*13*). Male and female *Ube3a-1x* mice were crossed to result in litters containing WT, *Ube3a-1x* and *Ube3a-2x* mice. *Ube3a-1x* mice express similar quantities of *Ube3a* mRNA but encod an untargeted UBE3A protein, comparing with *Ube3aNLS* mice which increases of UBE3A targeted to the nucleus through a C-terminal nuclear localization signal fusion engineered into the full length *Ube3a* gene. *Ube3a-2x* mice are homozygous for the C-terminal FLAG-tagged, full-length *Ube3a* transgene. To determine transgenic *Ube3a* copy number, genomic copies of *Ube3a* and *Lgi1* were quantified using real time PCR (*Ube3a* primers F: TACTGCTGAAGGTTTTCTTGGG, R: CTGCGAAATGCCTTGAATTGTT, Lgi1 primers F: ACCTAAGAGGGAACGCATTT, R: AATGATACAGTCAAAATCCT). KAPA SYBR FAST qPCR Master Mix (Kapa Biosystems) was used in 6*μ*l triplicate reactions containing 2.5 ng genomic DNA and 200 nM primers. Absolute *Ube3a* copy numbers were calculated using the 2^^-(^**^ΔΔC^t^)^**method, multiplying by a factor of two (two *Lgi1* alleles). The following lines were bred into an FVB/NJ genetic background for 6 or more generations. *Grid1*KO mice were generously provided by Dr. Jian Zuo (St. Judes Childrens Research Hospital, Memphis, USA). *Cbln1^flx/flx^* mice were a generous gift from Dr. Masayoshi Mishina (*24*) (Nigata University, Tokyo, Japan). *VGluT2-Cre* (*Slc17a6^tm2(cre)Lowl^*), *VGaT-Cre* (*Slc32a1^tm2(cre)Lowl^*), *Agrp-Cre* (*Agrp^tm1(cre)Lowl^*) mice and *Npy-hrGFP* (B6.FVB-Tg(Npy-hrGFP)1Lowl/J) mice were provided as a generous gift from Dr. Bradford Lowell (BIDMC). The *Sf1-Cre* (Tg(Nr5a1-cre)7Lowl)*, Pomc-Cre* (Tg(Pomc1- cre)16Lowl), *Tac1-Cre* (*Tac1^tm1.1(cre)Hze^*)*, Esr1-Cre* (*Esr1^tm1.1(cre)And^*)*, Pgr-Cre* (*Pgr^tm1.1(cre)Shah^*), and *ePet-Cre* (Tg(Fev-cre)1Esd) lines of mice, and *Nrxn1α* KO, Ai9 (RCL-tdT), Ai32/*ChR2-EYFP* (*34*) were purchased from Jackson Labs. To generate the conditional *Grid1* knockout FVB/NJ mice, an *Easi*-CRISPR strategy (*39*)was employed to insert a loxP recombination site within each of the introns flanking exon 4 of *Grid1 (NCBI ID: 14803).* Chemically-modified sgRNAs were obtained from Synthego; gRNA target 1: 5’-TCCAGCCCCAGGATATAAGG-3’, gRNA target 2: 5’-CGGTTCCTTCACAGACCACG-3.’ The single-stranded DNA homology-directed repair template containing the loxP-flanked *Grid1* exon 4 and homology arms was synthesized by Integrated DNA Technologies. CRISPR/Cas9-edited founder mice were generated at the BIDMC Transgenic Core Facility by microinjection of FVB/NJ zygotes with a mix of 200 ng/μl SpCas9- NLS, 50 ng/μl sgRNA1, 50 ng/μl sgRNA2 and 15 ng/μl ssDNA. Candidate founder mice were crossed to FVB/NJ to obtain germline-transmitted F1 mice. F1 mice with correct editing at the target locus were identified by sequence validation and used to establish the floxed *Grid1* mouse line.

### Viral vectors

AAV2-CMV-CreGFP, AAV2-CMV-GFP, AAV2-hSyn-DIO-GFP, AAV2-hSyn-DIO-hM3D(Gq)-mCherry, AAV2-hSyn-DIO-hM4D(Gi)-mCherry and AAV2-EF1α-DIO- ChR2(E123T/T159C)-EYFP were purchased from the University of North Carolina Viral Vector Core. AAV2-hSyn-DIO-Cbln1, AAV2/9-hSyn-DIO-Ube3a, AAV2/9-hSyn-mVglut2-mCherry and AAV2/9-hSyn-mVglut2-mCherry-2A-Cre were generated as previously described (*13*). AAV2/9-hSyn-DIO-Ube3a-NLS was generated by amplifying a *3xFLAG-2xNLS* cassette from *1xUbe3aNLS* genomic DNA using primers: 5’- TCTTCCGCATGCTGGACTATAAAGACCATGACGGTGAT-3’ 5’-TCTTCCGGATCCGGCGCGCCTTAGACCTTACGCTTCTTCTTAGGAC-3’ and subcloning into SphI and BamHI sites of the *pLVX-Ube3a-IRES-mCherry* construct. The resulting *Ube3a-NLS* cassette was then digested with SpeI and AscI and subcloned into NheI and AscI sites of the pAAV-hSyn-DIO-EGFP construct (Addgene, Plasmid #50457). pAAV-hSyn-DIO-Ube3a- NLS was then sent to the University of North Carolina Vector Core for AAV particle production with serotype 9. To generate the pAAV-mMeCP2-DIO-EGFP-W3SL vector, first, the W3SL enhancer sequence was amplified from Addgene Plasmid #61463 with a 5’ XhoI site addition and subcloned into EcoRI and KpnI sites of Addgene Plasmid #61591 to replace bGHpA. Next, the mMeCP2 promoter sequence was amplified from Addgene Plasmid #60957 with a 3’ SalI site addition and subcloned into XbaI and SpeI sites. Then, the DIO-EGFP cassette was amplified from Addgene Plasmid #50457 and subcloned into SalI and EcoRI sites: pAAV-mMeCP2-DIO-EGFP- W3SL. To generate Cre-conditional shRNA AAV constructs, the U6-TATAlox-CMV-EGFP- TATAlox cassette was amplified from pSico (Addgene Plasmid #11578) and subcloned into XbaI and XhoI sites of pAAV-mMeCP2-DIO-EGFP-W3SL to replace mMeCP2-DIO-EGFP. An HpaI site within W3SL was destroyed using the Q5® Site-Directed Mutagenesis Kit (NEB): pAAV-U6-TATAlox-CMV-EGFP-TATAlox-W3SL. For *Grid1* knockdown, three *Grid1* and one scrambled control shRNA hairpin sequences using TCTGCTT loop sequence were subcloned individually into HpaI and XhoI sites of pAAV-U6-TATAlox-CMV-EGFP-TATAlox-W3SL. shRNA target sequences were selected from the Broad Institute RNAi Consortium database: Grid1-A (GCTGAGAATATCCTTGGACAA) Grid1-B (GTGCTCATATTCGTGTTGAAT) Grid1-C (CGTTACAAAGGGTTCTCCATA) and Scrambled (CCTAAGGTTAAGTCGCCCTCG).

AAV-mMeCP2-DIO-EGFP-W3SL and AAV-U6-TATAlox-CMV-EGFP-TATAlox- Scrambled -shRNA W3SL vectors were packaged using AAV9 serotype at Boston Children’s Hospital Viral Core. For the AAV2/9-U6-TATAlox-CMV-EGFP-TATAlox-Grid1-shRNA-W3SL virus, the three different *Grid1* shRNA target plasmids were combined in equimolar ratio and packaged together in a single preparation. The AAV-hSyn-DIO-GCaMP7f-WPRE (AAV9 virus) are from Addgene.

### Stereotactic surgery

Anesthesia was induced in a chamber with isofluorane/oxygen following which mice were placed into a stereotaxic frame fitted with a continuous isofluorane delivery system. A single midline vertical scalp incision revealed skull landmarks. Stereotactic measurements were used to make 0.7mm wide burr holes over the entry point for bilateral viral injections directed at the VMHvl (0 deg, AP -1.5 mm, ML +/- 0.78 mm, DV -5.76 mm), VMHdm (0 deg, AP -1.46 mm, ML +/- 0.70 mm, DV -5.48 mm) and arcuate nucleus (0 deg, AP -1.7 mm, ML +/- 0.25 mm, DV -5.85 mm), all relative to bregma. 1 μl (or 0.5 μl for AAV2-EF1α-DIO-ChR2(E123T/T159C)-EYFP) of virus was infused at 0.2 μl/min through a 33 gauge Hamilton needle connected to an automated infusion pump. Following each infusion, the needle remained in place for 5 minutes. The incision was sutured closed and a single injection of 10 mg/kg meloxicam dissolved in saline was administered intraperitoneally for perioperative analgesia. All behavioral measurements were conducted approximately 28 days following surgery, and accurate targeting to the VMHvl and arcuate was confirmed by immunohistochemistry. Electrophysiological recordings were conducted at least 30 days after the surgery. The accuracy of injection sites at VMHvl and arcuate were also confirmed under microscopy when patch clamp recordings were performed.

### *In vivo* Ca^2+^ imaging with inscopix system

To prepare animals for in vivo Ca^2+^ imaging experiments in *Tac1-Cre* male mouse, a craniotomy was performed as described in stereotactic surgery above, and saline was repeatedly applied to the exposed tissue to prevent drying. The dura was then carefully removed with a 30-gauge beveled syringe needle. AAV-hSyn-DIO-GCaMP7f or AAV-hSyn-DIO-GCaMP7f + AAV-hSyn-DIO- Ube3a-NLS (300 nL) virus were injected unilaterally into VMHvl of *Tac1-Cre* male mice as described in stereotactic surgery procedures at the coordinates as (0 deg, AP -1.5 mm, ML +/- 0.78 mm, DV -5.76 mm). After finishing the virus injection, a ProView™ Lens Probe (0.5 mm diameter, 8.4 mm length, Inscopix, Palo Alto, CA) was implanted above the virus injection site with coordinates (−1.5 mm posterior to bregma, 0.6 mm lateral to midline, and −5.5 mm ventral to skull surface). The bottom of the lens was situated approximately 200 to 300 μm directly above the VMHvl. The portion of the lens extending ∼2 to 3 mm above the skull surface was fixed with dental cement (C&B Metabond® Quick Adhesive Cement System, Parkell Inc). A silicone elastomer (Kwik-Cast; World Precision Instruments, Sarasota, FL) was applied to the top of the lens to protect the imaging surface from external damage. Four to six week later after virus injection and lens implantation, baseplate (Inscopix, Palo Alto, CA) was installed to support the miniaturized microscope. For this procedure, mice were anesthetized with isoflurane, and the silicone mold around the lens was carefully detached. Debris was removed from the exposed lens with compressed air canisters, and lens paper with ddH2O was used to clean the top of the lens. Next, the miniature microscope (nVista 3, 475-nm blue LED; Inscopix, Palo Alto, CA) with the baseplate attached was positioned above the implanted lens with an adjustable gripper (Inscopix, Palo Alto, CA). The microscope was lowered toward the top of the lens by micromanipulator until the field of view was in focus. To ensure that the objective lens was completely parallel and aligned with the implanted lens, the angle of the microscope’s position was adjusted by manually tilting the scope within the adjustable gripper. Once the field of view was in focus and the scope was parallel with the lens, the magnetic baseplate was cemented around the implanted lens. Finally, a baseplate cover (Inscopix, Palo Alto, CA) was secured into the baseplate with a set screw to protect the lens until imaging. One week later after installation of baseplate, a female mouse was added to the cage of the *Tac1-Cre* male mouse for one week for preparing resident intruder (R-I) test. Then the *in vivo* Ca2+ imaging was performed in freely moving animal when R-I tests were conducted. For each imaging session, the nVista3 microscope was connected to the baseplate on the cranium and fixed in place by the baseplate set screw. Animals acclimate in their home cage for 20 min prior to the starting imaging sessions. Calcium images were acquired using Inscopix nVista3 Data Acquisition Software v1.3.1 with collecting frame rate 10 Hz with an average exposure time of 65 ms. The analog gain on the image sensor was set to 6, while the LED power was set as 1 mW. The acquired data by the Inscopix microscope were analyzed using Inscopix Data Processing Software (IDPS version 1.6) to visualize calcium spike movies, extract calcium traces, threshold events and calculate calcium spike frequencies. The *ΔF/F0 = [F(t) − F0]/F0*, where the baseline image (*F0*) was calculated using mean frame of the input movie, and calcium imaging data were normalized as relative changes in fluorescence. The cell images and traces were identified by running PCA/ICA algorithm or manual ROI. The calcium spikes were calculated after deconvolution of cell traces. The amplitude value of the calcium spikes was export as excel file from the IDPS. The calcium spike score (sum of amplitude values of calcium spikes during the first 5 minutes male exposure epoch) was used to comparing calcium events in resident *Tac1-Cre* male mice expressing AAV-DIO-GCaMP7f + AAV-DIO-Ube3a-NLS to mice only injected AAV-DIO-GCaMP7f.

### Behavioral measurements

Resident intruder paradigm (*40*): mature male mice (8-12 weeks old) were housed with a virgin FVB female (6-8 weeks old) for seven days in a room with an inverted light-dark cycle (The dark cycle began at 1:00 PM). Mice were transferred to the light/dark shifted room after weaning and acclimated for a minimum of two weeks. On the testing day the female mouse was removed at 10:00 AM and the first test was conducted at 1:30 PM. WT C57BL/6 male mice were used as intruder mice that were group-housed (five per cage) and matched with resident mice for approximate age (8 weeks old) and body weight (weighing less than the resident mice). The resident’s home cage was placed into a fume hood, and an intruder mouse was introduced into the resident’s cage, then their interactions were video recorded for 10 minutes. Aggressive behavior was video recorded and also scored live by a trained viewer blinded to genotypes. For studies involving DREADDs (designer receptors exclusively activated by designer drugs) (*41*) (i.e. AAV2-DIO-hM3D(Gq)-mCherry and AAV2-DIO-hM4D(Gi)-mCherry), approximately 28 days following AAV injections, mice received intraperitoneal injections of clozapine-N-oxide (CNO, Sigma, dissolved in saline) 10 minutes prior to the resident-intruder test.

For *in vivo* Ca^2+^ imaging with inscopix during resident-intruder test, aggression behavior was recorded with a Basler color camera, which was controlled by the Ethovision XT 15 software (Noldus). Inscopix Ca2+ imaging was synchronized with the behavior recordings through a USB I/O box which also controlled by Ethovision XT 15. Inscopix Data Processing Software was used to analyze the Ca2+ imaging data. Synchronous video recordings were used to manually score aggression behaviors.

### Immunofluorescence and c-fos quantifications

Mice were anesthetized with an intraperitoneal injection of tribromoethanol and then perfused with ice-cold PBS followed by cold 4% paraformaldehyde. Brains were frozen in optimal cutting temperature (OCT) compound (Fisher Health Care) after cryoprotected in 15% sucrose followed by 30% sucrose (each for 24h). 16µm sections were cut on a cryostat and slide-mounted. When an anti-mouse secondary antibody was used, sections were blocked with MOM reagent (Vector) and then sections were permeabilized with 0.1% Triton X100 and blocked with 10% normal goat serum with 1% BSA in PBS. The FLAG tag, GFP and mCherry were probed respectively with FITC-conjugated anti-FLAG (1:200, Sigma), Green Fluorescent Protein (GFP) antibody (JL-8, 1:500, Clontech), Red Fluorescent Protein Antibody (DsRed, 1:500, Clontech). The brain sections were incubated with antibodies in blocking solution at 4°C temperature overnight. Sections were washed and when required, incubated with Alexa-conjugated secondary antibodies (1:500, Invitrogen) for 3 hours at room temperature, and then mounted in Vectashield with DAPI (Vector).

Fluorescent images were taken using a LSM510 confocal microscope (Zeiss), FV1000 Confocal microscope (Olympus) or the VS120-SL 5 Slide Scanner (Olympus). Images were processed using Fiji image J software.

For c-fos quantifications (Fig. 3, Fig S4), animals were sacrificed 75 minutes after resident-intruder test, then perfused with 4% paraformaldehyde. Brains were frozen in OCT compound (Fisher Health Care) after cryoprotected in 15% sucrose followed by 30% sucrose (each for 24h). Coronal brain sections (50 µM, from bregma +3mm to -6mm) were cut on a cryostat. Sections were processed as described above for c-Fos antibody immunohistochemical staining (c-Fos Rabbit mAb #2250, 1:2000, Cell Signaling Technologies). Brain sections mounted on the slides were scanned using Zeiss Axio Scan.Z1 with 5x magnification. The whole slide images were sorted sequentially from rostral to caudal using image processing software QuPath (*42*) referenced to The Mouse Brain in Stereotaxic Coordinates Atlas (Paxinos and Franklin, Compact 3rd Edition). The GFP channel raw images were exported with the brain structure of interest and surrounding landmarks. Regions of Interest (ROIs) was selected on the exported GFP channel images using Fiji ImageJ software. ROIs were saved and applied for same-bregma slides in different mouse sections with minor adjustment for individual differences. A Macro including ImageJ functions of Brightness/Contrast adjust, Background Subtract, Median Filter and Auto Threshold (Yen module) was used to threshold images in each ROI consistently. The c-fos positive neurons were detected with StarDist 2D plugin after threshold, then converted to binary images for c-fos counting with Analyze Particle function using ImageJ. The masked images highlighting counted c-fos positive neurons within each ROI then be shown on raw images (fig. S4, D and F).

### Tissue preparation, RNA isolation and quantitative real time PCR (qRT-PCR)

Brains were rapidly removed and submerged in ice-cold phosphate buffered saline (PBS) for 20- 30s. For the isolation of cortex (ChIP studies), two “slabs” of total cortex were dissected using curved forceps. For VMH punches, the PBS-cooled brains from wild type and *Ube3aNLS7-1x* mice (n=6 each) were placed in a brain matrix (ASI Instruments) and razor blades were placed at 1mm intervals to obtain brain slices containing VMH. Punches were obtained with a 0.5mm punch core needle (Harris Unicore) and immediately frozen to -20C and stored at -80C. RNA was isolated using Trizol Reagent (ThermoFisher) and column-purified using the RNAeasy Protect Mini Kit (Qiagen). First strand cDNA synthesis was carried out with M-MLV Reverse Transcriptase (ThermoFisher) and Oligo (dT) 20. Expression of *Cbln1* was quantified using RT-qPCR with Power SYBR Green Master Mix (ThermoFisher) and a Bio-Rad CFX 384 Real-Time System. Primer pairs were selected through Primer3 with specificity confirmed with Primer-BLAST. Primer sequences are listed in Supplementary Table2. *Cbln1* expression between genotypes was determined using the ΔΔCT method with *Syn1* as a reference gene. Melt-curve analysis and agarose gel electrophoresis were used to confirm a single PCR product of the appropriate amplicon size.

### Luciferase Assay

The *Cbln1* promoter luciferase construct was purchased from GeneCopoeia, Inc. (Cat# MPRM22597-PG02), which encodes Gaussia luciferase driven by a 1331 bp genomic fragment upstream of the coding sequence of the murine *Cbln1* gene (Accession: NC_000074.6). The *Ube3a* expression construct was generated by amplifying the coding sequence of human Ube3a isoform III from Plasmid #37605 (Addgene) with primers *5’-TCTTCCACTAGTGCCACCATGGCCACAGCTTGTAAAAGATC-3’* and *5’-TCTTCCGGATCCTTACAGCATGCCAAATCCTTTGG-3’* and subcloning into the SpeI and BamHI sites of the *pLVX-IRES-mCherry* vector (Clontech Cat#631237). HEK293T cells were transfected in 96-well using Lipofectamine 3000 Reagent (ThermoFisher) with 50 ng *Cbln1* promoter-Gluc plasmid and 50 ng of either *pLVX-IRES-mCherry* or *pLVX-Ube3a-IRES-mCherry* per well (*n* = 6 each condition). 48 hours after transfection, Gaussia luciferase activity was developed using the Secrete-Pair Gaussia Luciferase Assay Kit (Genecopoeia, Inc., Cat#LF061) and measured with the BioTek Synergy 2 luminescence plate reader.

### Chromatin immunoprecipitation

25 mg cortical tissues were dissected from wild type mice (*n* = 3) and chromatin was prepared using the SimpleChIP plus Enzymatic Chromatin IP Kit (Cell Signaling). UBE3A-bound chromatin was immunoprecipitated using rabbit anti-UBE3A antibody (Bethyl Laboratories). Rabbit IgG was used as a mock control. The ratio of UBE3A-bound *Cbln1* promoter DNA relative to input chromatin was quantified using qRT-PCR with primers *5’- CTCGCCGCTCCTAATAACAA-3’, 5’- CCACCCTCCAGCCAATC-3’* and FAM/ZEN-labeled probe *5’-ACAGGGCAACCATTGGCTCG-3’* with Taqman Gene Expression Master Mix (ThermoFisher).

### Brain slice patch clamp electrophysiology and *in vitro* optogenetics

Mice were deeply anesthetized with isofluorane, following which mouse brains were rapidly removed and submerged in an ice-cold artificial cerebrospinal fluid (ACSF) containing (in mM): 126 choline chloride, 2.5 KCl, 1.2 NaH2PO4, 1.3 MgCl2, 8 MgSO4, 0.2 CaCl2, 20 glucose, and 46 NaHCO3, equilibrated with 95% O2/5% CO2. Coronal brain slices containing the VMHvl or arcuate nucleus (300 μm thick) were made using a Leica VT1200S microtome (Leica Microsystems). The slices were transferred into a holding chamber containing ACSF comprised of (in mM): 124 NaCl, 3 KCl, 1 MgSO4, 1.25 NaH2PO4, 2 CaCl2, 25 Glucose, and 26 NaHCO3, equilibrated with 95% O2/5% CO2. The slices were maintained at 33°C temperature for 30 minutes and then kept at room temperature until recorded. To increase viability of neurons, 5 μM glutathione, 500 μM pyruvate, and 250 μM kynurenic acid were added to the choline chloride replacement ACSF and ACSF in the holding chamber. During recordings, slices were perfused (2 ml/min) with ACSF without glutathione, pyruvate and kynurenic acid. Whole cell recordings were performed from neurons located in the VMHvl or arcuate nucleus under visual control on an upright microscope with differential interference contrast and infrared illumination. Arcuate NPY neurons were identified by GFP fluorescence labeled by *NPY-hrGFP* allele. Patch pipettes were pulled using a P-97 puller (Sutter Instruments) and filled with (mM): 130 K gluconate, 10 HEPES, 5 NaCl, 1 MgCl2, 0.02 EGTA, 2 MgATP, 0.5 NaGTP and 10 mM sodium phosphocreatine at a pH of 7.3 and osmolality of 275 to 290 mOsm. Filled patch pipettes had resistances of 3-5 MΩ. Synaptic currents were recorded in voltage clamp mode by an Axon MultiClamp 700B amplifier (Molecular Devices). Data were filtered at 1 kHz, digitized at 10 kHz with DigiData 1440A interface (Molecular Devices) and acquired by Clampex 10.5. Series resistance was monitored with a 5 mV hyperpolarizing step (5 ms). Resting membrane potential and action potentials (APs) were recorded in current clamp mode with APs induced by injecting 200 ms square wave pulse of positive currents ranging from 40 to 400 pA. An optical fiber (200 μm core diameter, 0.2 numerical apertures) was used to deliver optical stimulation. The fiber was placed 200 μm from the site of recording and coupled to a diode-pumped 473 nm laser (CrystaLaser Company) to control stimulus intensity light pulse duration (5ms) was controlled by a Master-8 stimulator and light pulse frequency (0.1 Hz) was determined and triggered by the Clampex 10.5. The evoked EPSCs in neurons-expressing channelrhodpsin-2 (ChR2) were induced by the laser light (30 mW)(*33*). 12 consecutive sweeps were averaged with Clampfit 10.5 to determine the light-evoked EPSC amplitude. GABAA antagonist bicuculline (10 μM) and NMDA receptor antagonist D-APV (30 μM) were present to isolated the AMPA receptor currents. 30 μM 6-cyano-7-nitroquinoxaline-2,3- dione (CNQX) was used in some studies to block and confirm AMPA receptor currents. Miniature EPSCs (mEPSCs) were recorded at a holding potential of -70 mV in the presence of GABA_A_ antagonist bicuculline (10 μM) and tetrodotoxin (TTX, 300 nM). mEPSCs were detected and analyzed using Mini Analysis Program (Synaptosoft, Decatur, GA). Amplitude and area thresholds were set to 5 pA and 20 fC, respectively. The peak amplitude and inter-event interval of mEPSCs from 60 second episodes were used to generate cumulative probability plots, and the statistical significance was determined by Kolmogorov-Smirnov test.

### Single neuron RT-qPCR in VMHvl

The brain slices were prepared as described in electrophysiological recordings. Then single tdTomato fluorescent labeled neuron (Ai9 mice) were aspirated from the VMHvl and collected into 5 μl of 2x CellsDirect Reaction Mix (ThermoFisher, Cat# 11753-100). cDNA was generated and preamplifed 20 cycles for target sequences and treated with ExoSAP-IT (Affymetrix, Cat# 78201) as previously described (*43*). Expression of *Cbln1* and other target genes was quantified using RT-qPCR with Power SYBR Green Master Mix (ThermoFisher) and a Bio-Rad CFX 384 Real-Time System. Primer sequences are listed in Supplementary Table 2. Fold change in *Cbln1* expression between genotypes was determined using the ΔΔCT method with *Syn1* as a reference gene.

### Statistics

Sample sizes are shown in figure legends in parentheses and were chosen based on a power analysis designed to exceed a statistical power (1-beta) of 80% and to meet or exceed sample sizes typically employed in similar mouse behavioral experiments. Behavioral studies performed in a blinded fashion and the code broken after the testing when data was analyzed. Where possible, animals were placed into groups through a randomization process. No animal are excluded from the data analysis. For resident–intruder aggression tests, an unpaired two-tailed Student’s t-test was used to determine statistical significance when comparing two groups. Comparisons across two groups before and after CNO treatment were analyzed by paired two-tailed Student’s t-test. Multiple groups were analyzed using 1-way ANOVA followed by Bonferroni’s Multiple Comparison Correction. 2-way Repeated-Measures ANOVA followed by Multiple Comparison was used to analyze c-fos quantification data (Fig. 3C-F and fig. S4, B, C, E and F). One-tailed Fisher’s exact test was used to analyze the data of male attacking female in (fig. S3C). The mEPSC amplitudes and inter-event intervals from 60 second episodes were used to generate cumulative probability plots, and the statistical significance was determined by Kolmogorov-Smirnov test (KS). Data are presented as mean ± standard error of the mean (SEM). *P* < 0.05 was considered statistically significant with *ns* indicating non-significant, \**P* <0.05, \*\**P* < 0.01, \*\*\**P* <0.001 and \*\*\*\**P* <0.0001.

### Materials Availability

Mice created for this manuscript have been deposited at JAXS.

## Acknowledgements

We thank Anderson Lab members Oriana DiStefano, Greg Salimando, Rebecca Broadhurst and Scott Rochard for mouse colony upkeep and genotyping and Vaishnav Krishnan and Marcello DiStasio for reading and editing the manuscript. We thank the Harvard Center for Biological Imaging at Harvard University, the Neurobiology Imaging Facility at Neurobiology Department of Harvard Medical School for consultation and instrument availability that supported this work (in part through a NINDS P30 Core Center grant #NS07203) and Boston Children’s Hospital IDDRC (1U54HD090255, P30HD18655). This work was supported by funding to M.P.A. from The National Institute of Mental Health (R01MH112714, R01MH114858, and 1R21MH100868), The National Institute of Neurological Disorders and Stroke (1R01NS08916), The Eunice Kennedy Shriver National Institute of Child Health and Human Development (1R21HD079249), The Nancy Lurie Marks Family Foundation, Landreth Foundation, Autism Speaks/National Alliance for Autism Research, and the Simons Foundation.

## Author Contributions

Y.N., D.C.S., M.A.J., M.B., and M.P.A. designed the study. Y.N., D.C.S., M.B., and M.P.A wrote the manuscript. Y.N, D.C.S., M.B., and. M.A.J., performed all the experiments and analyses except for the following: J.T. performed some aggression experiments; J.S. and X.Z. performed some data analysis; C.R. and Y.H. performed some immunostaining; M.N., I.N., and E.M.K. developed molecular probes. Authorship order amongst the starred first authors was determined by the relative contribution to total data, technologies, and writing. Competing Financial Interests: Y.H., Y.N., and M.P.A became employees of Regeneron Pharmaceutical.

Supplementary Materials: the list of the supplementary materials:

Materials and Methods figures S1-6

## SUPPLEMENTARY FIGURES

**fig. S1.**
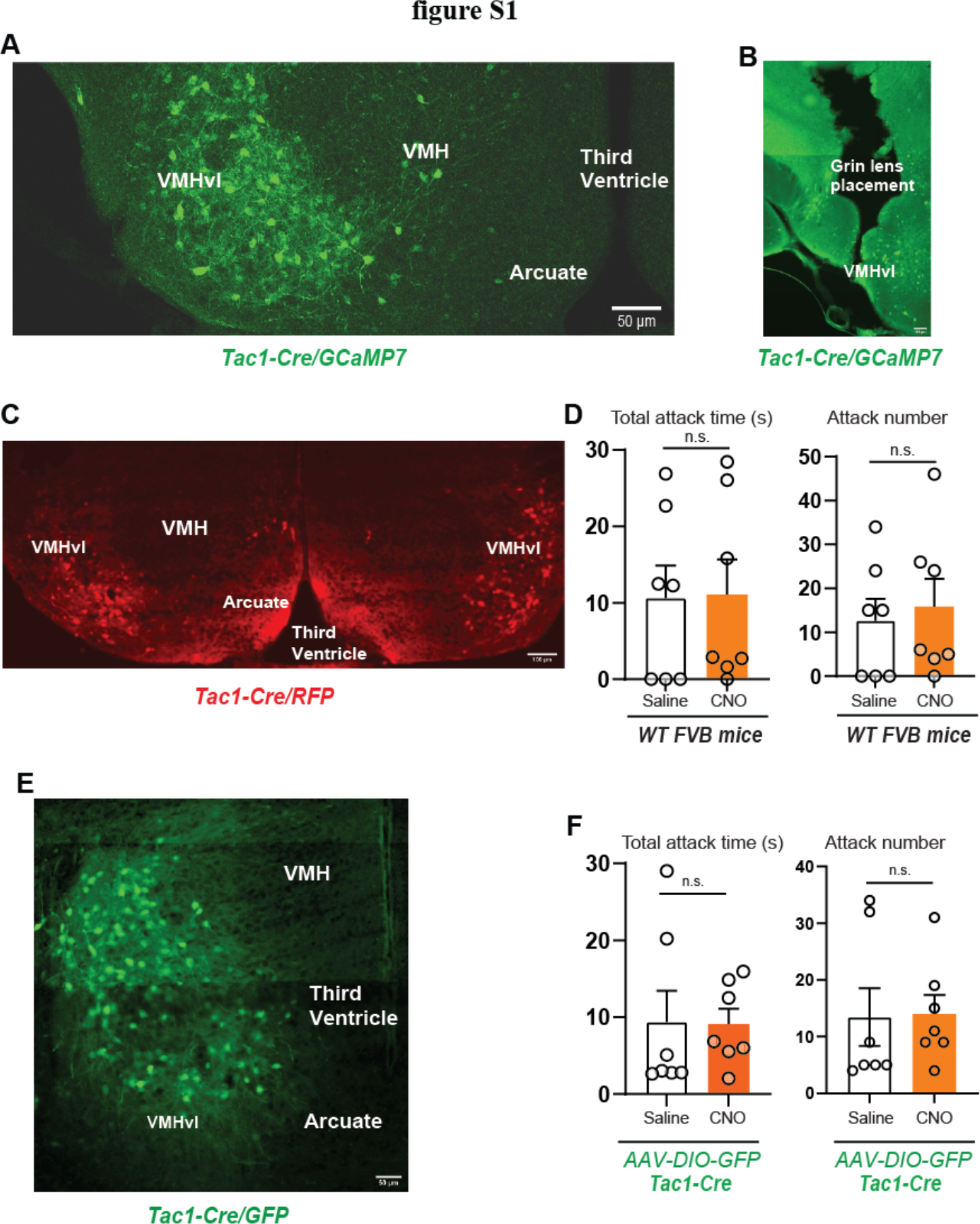
VMHvl *Tac1* neurons drive attack behavior. (**A**) Representative image of *GCaMP7f* expression in VMHvl *Tac1* neurons after stereotactic injection of AAV9-hSyn-DIO-GCaMP7f virus into VMHvl of *Tac1-Cre* male mice (scale bar, 50 μm). (**B**) Representative image of Grin lens placement in VMHvl of AAV9-hSyn-DIO- GCaMP7f injected *Tac1-Cre* male mice (scale bar, 100 μm). (**C**) Representative image of anti-DsRed antibody immunofluorescence staining in *Tac1-Cre* mice injected with AAV-DIO-hM3D(Gq)-RFP in VMHvl (scale bar, 100 μm). (**D**) Total attack time/number in wild-type *FVB* mice comparing application of saline with CNO (1 mg/kg i.p., *n* = 7, *P_T_* = 0.9429, *P_N_* = 0.6924). (**E**) Representative image of *GFP* expression in VMHvl *Tac1* neurons after stereotactic injection of AAV-DIO-GFP virus into VMHvl of *Tac1-Cre* male mice (scale bar, 50 μm). (**F**) Total attack time/number in *Tac1-Cre* male mice comparing application of saline with CNO (1 mg/kg i.p., *n* = 7, *P_T_* = 0.9564, *P_N_* = 0.9269). *P* < 0.05 was considered statistically significant with *ns* indicating non-significant, \**P* <0.05, \*\**P* < 0.01, \*\*\**P* <0.001 and \*\*\*\**P* <0.0001.

**fig. S2.**
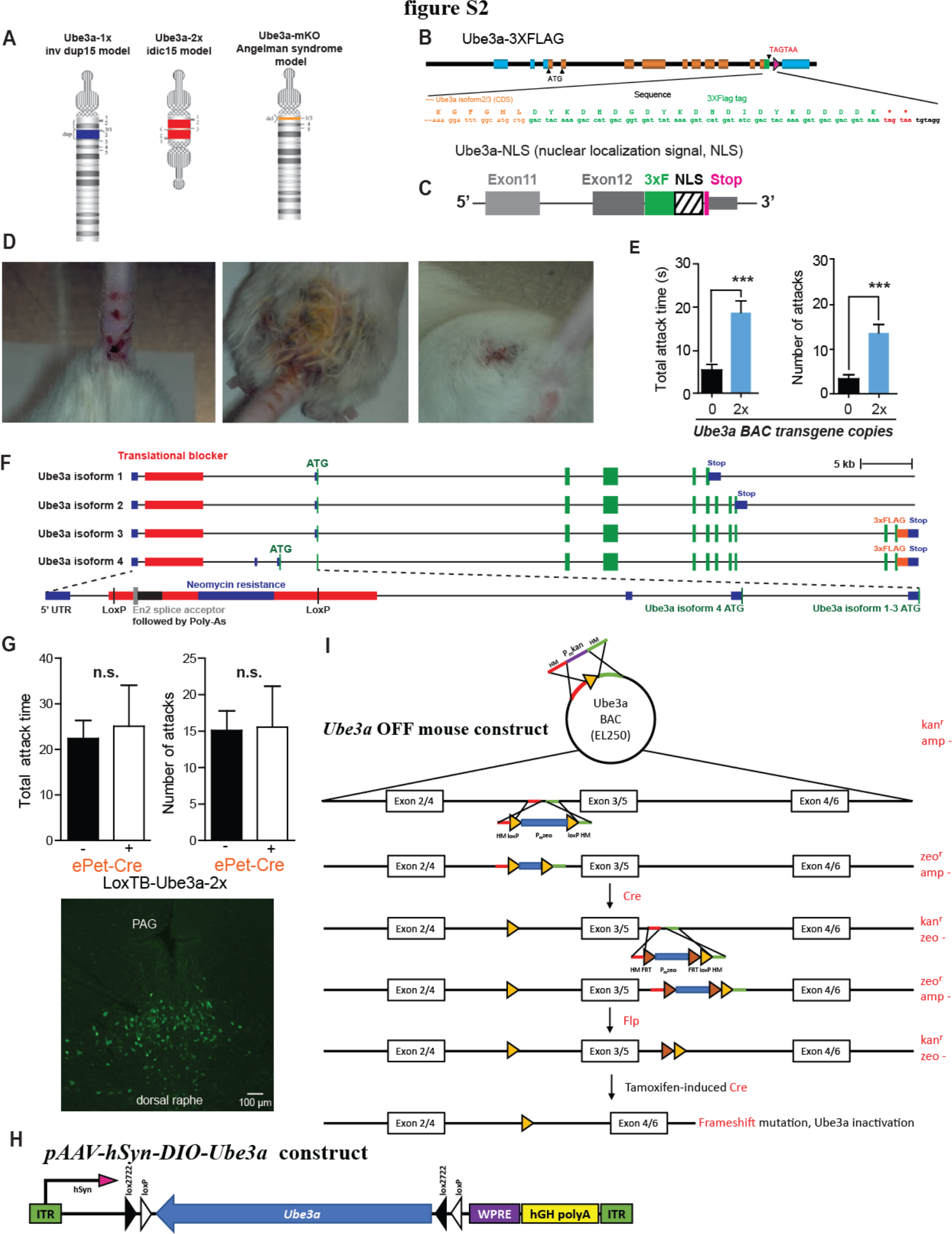
*Ube3a* genetic and molecular constructs. **(A**) Diagram of chromosome abnormality of maternal 15q11-13 interstitial duplication, maternal extranumerary isodicentric chromosome 15 (idic 15) and Angelman syndrome. (**B**) Diagram of construct of *Ube3a* 3xFlag transgenic mice with extra gene copies of full-length *Ube3a* gene. (**C**) Diagram of Ube3aNLS with 3xFLAG and nuclear localization signal (NLS) followed by a STOP codon added in frame to exon 12 of mouse Ube3a gene. (**D**) Photos of injured mice from cages with *Ube3a-2x* male mice. (**E**) Total attack time/number in wild type (*n* =23) and *Ube3a-2x* mice (*n* = 9, *P_T_* = 0.0007 and *P_N_* = 0.0003). (**F**) Diagram of *LoxTB-Ube3a* construct with transcriptional/translational stop cassette (red) containing an *En2* splice acceptor (grey) followed by polyA tails (black), all flanked by *LoxP* sites and inserted into intron 1 of full-length FLAG- tagged *Ube3a* gene designed to block all potential splice isoforms. (**G**) Upper panel: total attack time/number in *LoxTB-Ube3a-2x* mice (*n* = 20) comparing to *ePet-Cre* (Tg(Fev-cre)1Esd):*LoxTB- Ube3a-2x* mice (*n* = 9) (*P_T_* = 0.7520, *P_N_* = 0.9340). Lower panel: FLAG immunofluorescence in dorsal raphe, scale bar 100 μm. (**H**) Diagram of construct of *AAV-hSyn-DIO-Ube3a* that expresses *Ube3a* in a Cre-dependent manner. (**I**) Diagram of construct of *Ube3a*OFF#5, a conditional *Ube3a* transgene where *LoxP* site flank exons of the full-length untagged *Ube3a* gene. *P* < 0.05 was considered statistically significant with *ns* indicating non-significant, \**P* <0.05, \*\**P* < 0.01, \*\*\**P* <0.001 and \*\*\*\**P* <0.0001.

**fig. S3.**
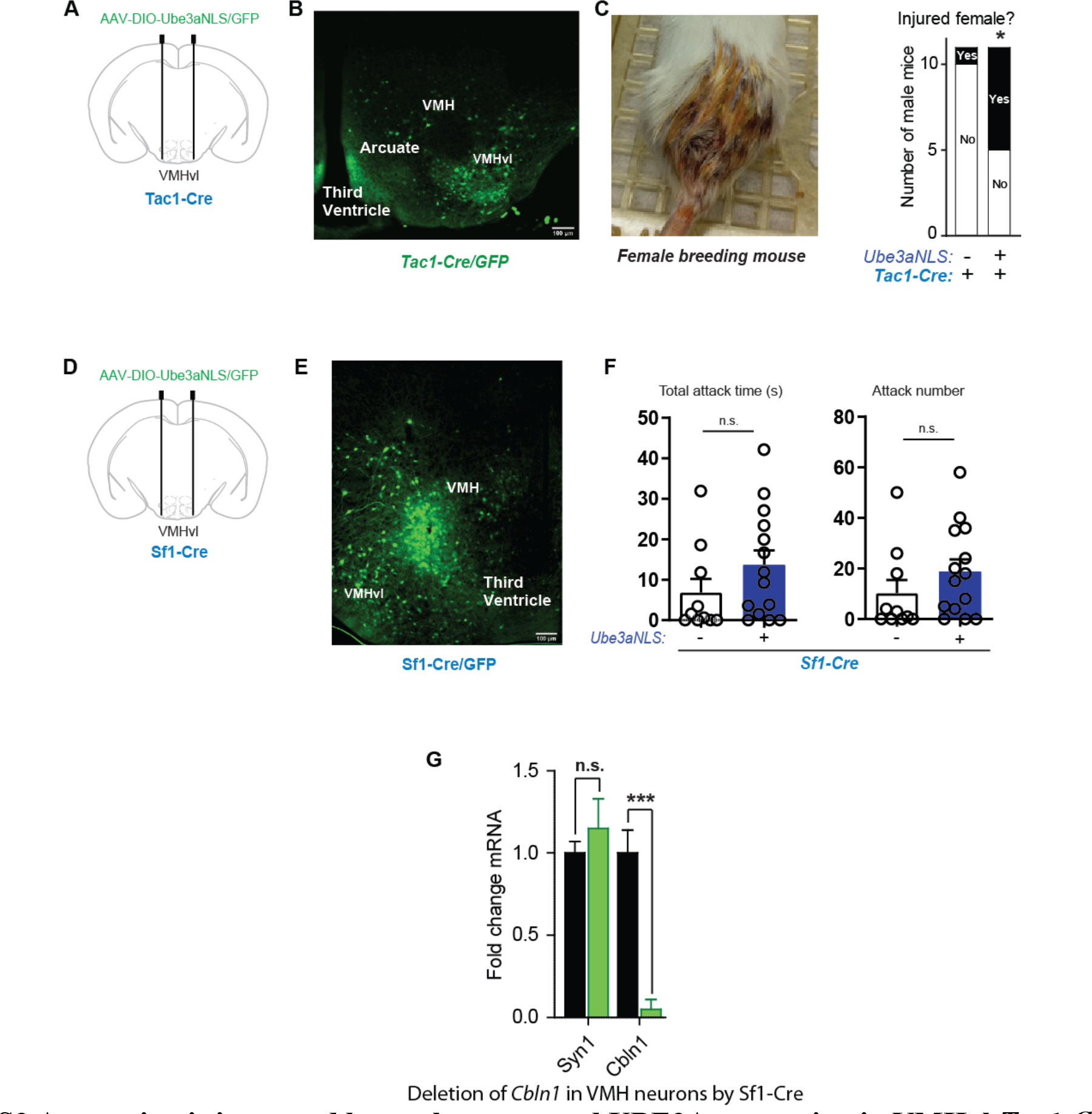
Aggression is increased by nuclear-targeted UBE3A expression in VMHvl *Tac1-Cre* neurons but not in VMHdm of *Sf1-Cre* neurons. (**A**) Diagram of stereotaxic injection. (**B**) Representative image of *GFP* expression in VMHvl *Tac1* neurons when stereotactically injecting AAV-DIO-Ube3a-NLS + AAV-DIO-GFP virus into VMHvl of *Tac1-Cre* male mice (scale bar, 100 μm). (**C**) Left: representative photo of female mouse with hind injury. Right: number of male mice that injured female added to their cages in *Tac1-Cre* mice injected with AAV-DIO-GFP + AAV-DIO-Ube3aNLS (*n* = 11 mice) compared to AAV-DIO-GFP (*n =* 11 mice) in VMHvl (*P* = 0.032). (**D**) Diagram of stereotaxic injection. (**E**) Representative image of *GFP* expression in VMH *Sf1* neurons when stereotactically injecting AAV-DIO-Ube3a-NLS+ AAV-DIO-GFP virus into VMHdm of *Sf1-Cre* male mice (scale bar, 100 μm). (**F**) Total attack time/number in *Sf1-Cre* male mice compared stereotactically injecting AAV-DIO-Ube3a-NLS+ AAV-DIO-GFP virus (*n* = 14) into VMHdm of *Sf1-Cre* male mice with only AAV-DIO-GFP (*n* = 10, *P_T_*= 0.2002, *P_N_* = 0.2460). (**G**) Combined data of quantitative RT-qPCR of *Cbln1*and *Syn1* mRNA from single neurons isolated from VMHvl in *Sf1-Cre*:*Cbln1^flx/flx^* (*n* = 24 neurons, green bar) comparing to *Cbln1^flx/flx^* (*n* = 28 neurons, black bar) (*P* = 0.0003). *P* < 0.05 was considered statistically significant with *ns* indicating non-significant, \**P* <0.05, \*\**P* < 0.01, \*\*\**P* <0.001 and \*\*\*\**P* <0.0001.

**fig. S4.**
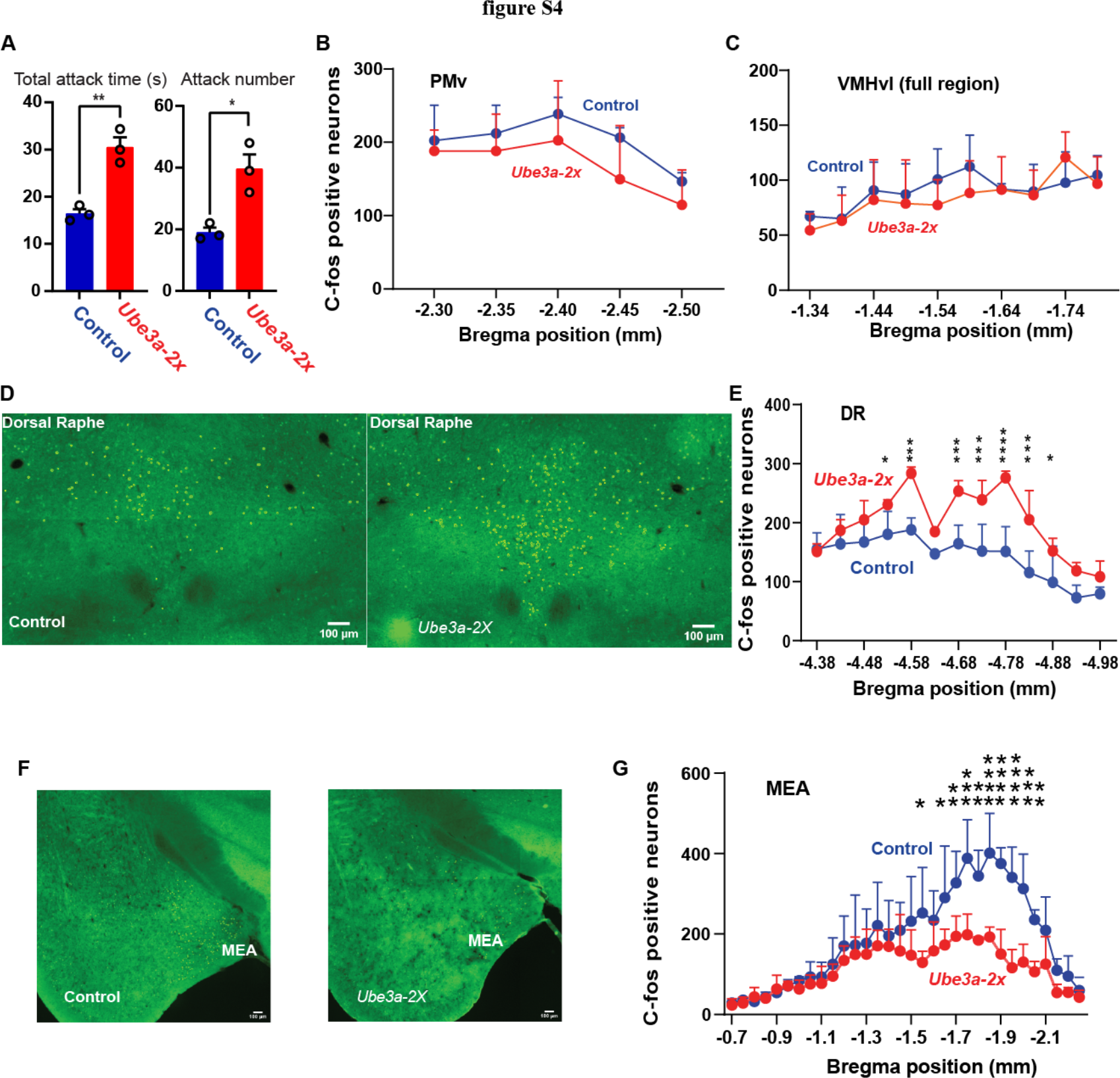
Images of anti-c-fos staining and quantification of c-fos positive neurons after aggression behavior test comparing *Ube3a-2x* with littermate control mice. (**A**) Total attack time/number in aggression behavior test compared *Ube3a-2x* (n = 3) and littermate control mice (n = 3, *P_T_* = 0.0035, *P_N_* = 0.0133). (**B**) and (**C**) Counts of c-fos positive neurons in ventral premamillary (PMv) (**B**, *P* = 0.3731) and full VMHvl (**C**, *P* = 0.3167) after aggression behavior test compared *Ube3a-2x* compared to control littermate mice. (**D**) Representative image of anti-c-fos antibody staining in dorsal raphe compared *Ube3a-2x* and control littermate mice (scale bar, 100 microns). (**E**) Counts of c-fos positive neurons in dorsal raphe (*P* = 0.0217) after aggression behavior testing comparing *Ube3a-2x* and control littermate mice. (**F**) Representative image of anti-c-fos antibody staining in MEA compared *Ube3a-2x* and control littermate mice (scale bar, 100 microns). (**G**) Counts of c-fos positive neurons in MEA (*P* = 0.1150) after aggression behavior test. All c-fos studies compared male *Ube3a-2x* (*n* = 3 mice) and control littermate mice (*n* = 3 mice). *P* < 0.05 was considered statistically significant with *ns* indicating non-significant, \**P* <0.05, \*\**P* < 0.01, \*\*\**P* <0.001 and \*\*\*\**P* <0.0001.

**fig. S5.**
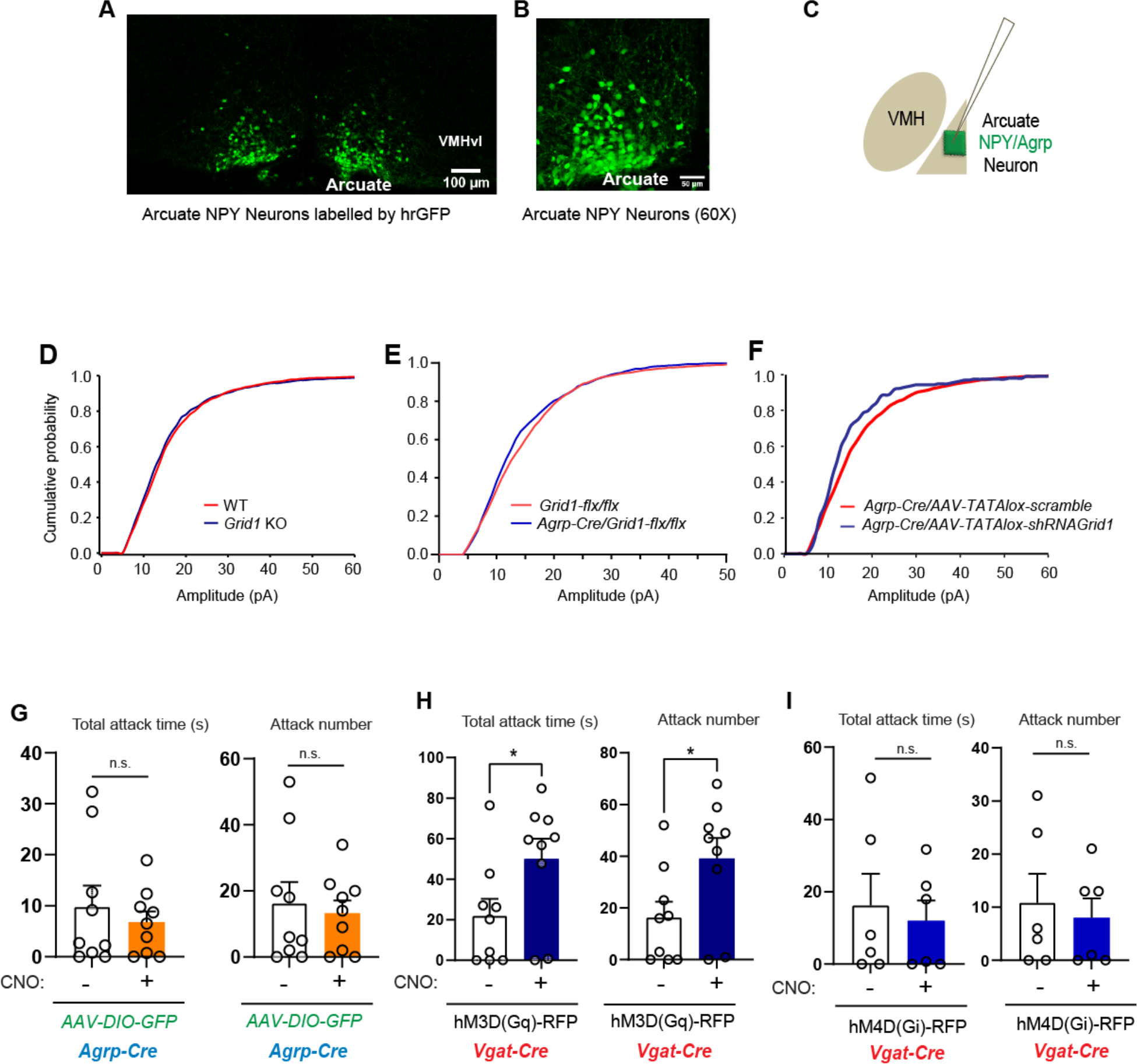
Miniature excitatory postsynaptic current (mEPSC) amplitude in arcuate NPY-GFP neurons when with *Grid1* loss of in arcuate AgRP/NPY neurons. (A) Representative image of GFP^+^ arcuate NPY neurons (labeled by *Npy-hrGFP* allele, scale bar, 100 microns) where whole– cell patch clamp recordings in arcuate were performed. (B) Higher magnification (60x) of GFP^+^ arcuate NPY neurons (scale bar, 50 microns). (C) Diagram of whole–cell patch-clamp recordings performed in GFP-labeled arcuate NPY/AgRP neurons. (D) Cumulative frequency plot of mEPSC amplitude in arcuate NPY GFP^+^ neurons in mice with homozygous deletion of *Grid1* (*n* = 4 mice, 16 cells) and control mice (*n* = 5 mice, 14 cells, KS test, *P* > 0.05). (**E**) Cumulative mEPSC amplitude in arcuate NPY GFP^+^ neurons in mice with *Grid1* knockout in arcuate AgRP/NPY neurons (*Grid1*-flx:flx/Agrp-Cre; *n* = 6 mice, 11 neurons) compared to control wild type mice (*n* = 4 mice, 14 cells, KS test, *P* > 0.05). (**F**) Cumulative mEPSC amplitude in arcuate NPY GFP^+^ neurons in mice with knockdown of *Grid1* in arcuate AgRP/NPY neurons (AAV- TATAlox-shRNA-Grid1 in Agrp-Cre; *n* = 3 mice, 6 neurons) compared to control wild type mice (*n* = 6 mice, 18 cells, KS test, *P* < 0.001). (**G**) Total attack time/number in *Agrp-Cre* mice (*n* = 9) injected with AAV-DIO-GFP comparing saline and CNO (1 mg/kg i.p., *P_T_* = 0.5232, *P_N_* = 0.6950). (**H**) Total attack time/number in *Vgat-Cre* mice (*n* =9) injected with AAV-DIO- hM4D(Gq)-RFP in the VMHvl comparing saline and CNO (1 mg/kg i.p., *P_T_* = 0.0477, *P_N_* = 0.0371). (**I**) Total attack time/number in *Vgat-Cre* mice (*n* = 6) injected with AAV-DIO- hM4D(Gq)-RFP in the VMHvl comparing saline and CNO (1 mg/kg i.p., *P_T_* = 0.6965, *P_N_* = 0.6738). *P* < 0.05 was considered statistically significant with *ns* indicating non-significant, \**P* <0.05, \*\**P* < 0.01, \*\*\**P* <0.001 and \*\*\*\**P* <0.0001.

**fig. S6.**
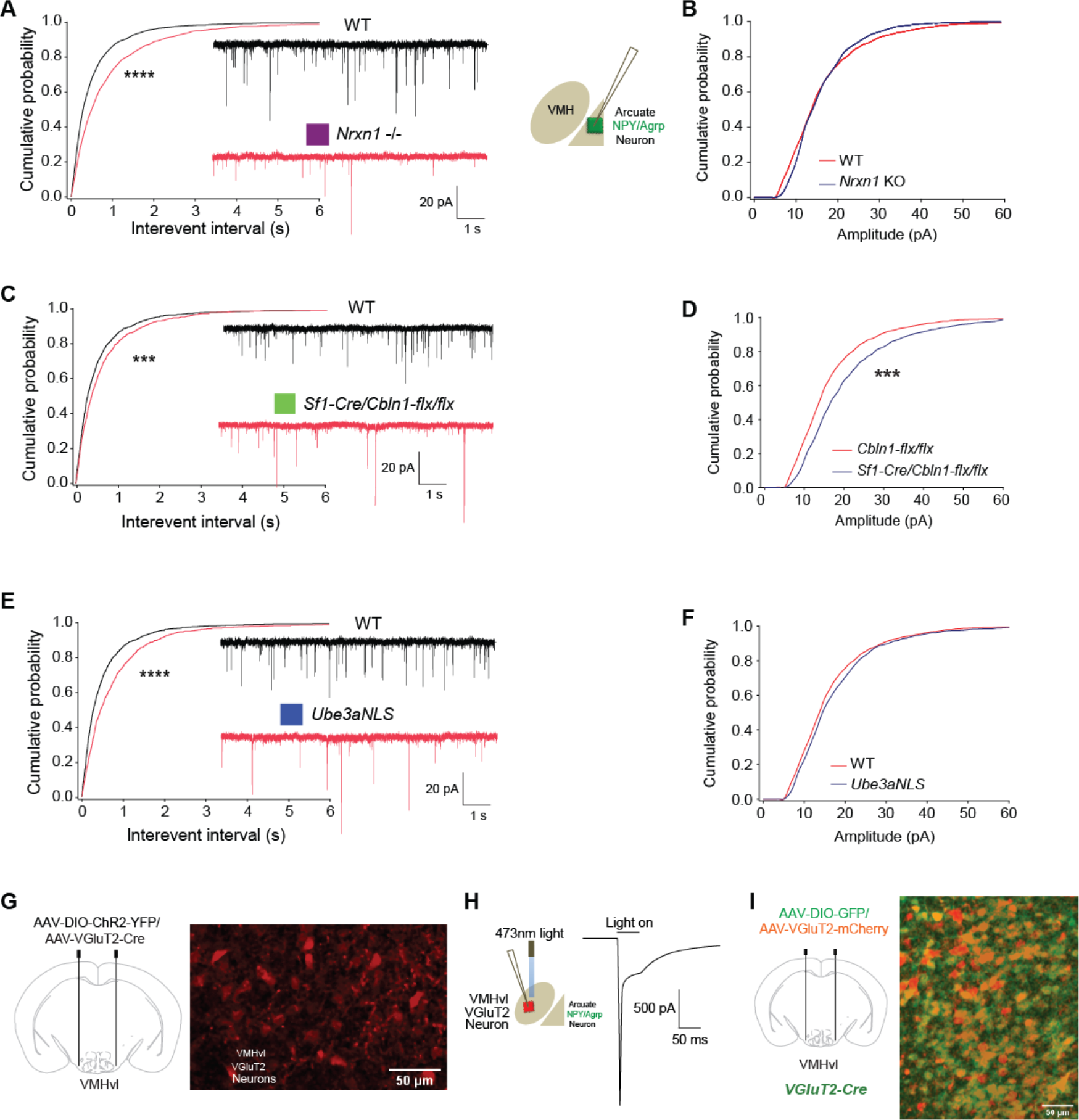
Miniature excitatory postsynaptic current (mEPSC) inter-event intervals and amplitude in arcuate NPY-GFP neurons of mice with deletion of *Nrxn1*, *Cbln1* or *Ube3aNLS7-1x*. (A) Left: Cumulative frequency plot of mEPSC inter-event intervals in arcuate NPY GFP^+^ neurons in mice with homozygous deletion of *Nrxn1* (*n* = 6 mice, 19 cells) and control mice (*n* = 5 mice, 14 cells, KS test, *P* < 0.00001); Right: diagram of whole–cell patch-clamp recordings were performed in GFP-labeled arcuate NPY/AgRP neurons. (**B**) Cumulative frequency plot of mEPSC amplitude in arcuate NPY GFP^+^ neurons (labeled by *NPY-hrGFP* allele) in mice with homozygous deletion of *Nrxn1* (*n* = 6 mice, 19 cells) and control mice (*n* = 5 mice, 14 cells, KS test, *P* > 0.05). (**C**) mEPSC inter-event intervals in arcuate NPY GFP^+^ neurons from *Sf1-Cre*:*Cbln1-flx/flx* mice (*n* = 5 mice, 19 cells) and control mice (*n* = 5 mice, 14 cells, KS test, *P* < 0.001). (**D**) Cumulative frequency plot of mEPSC amplitude in arcuate NPY GFP^+^ neurons in mice with *Sf1-Cre*:*Cbln1^flx/flx^* (*n* = 5 mice, 19 cells) and control mice (*n* = 5 mice, 14 cells, KS test, *P* < 0.001). (**E**) mEPSC inter-event intervals in arcuate NPY GFP^+^ neurons in mice with *Ube3a- NLS7*-*1x* (*n* = 5 mice, 17 cells) and control mice (*n* = 5 mice, 14 cells, KS test, *P* < 0.00001). (**F**) Cumulative frequency plot of mEPSC amplitude in arcuate NPY GFP^+^ neurons in mice with *Ube3aNLS7-1x* (*n* = 5 mice, 17 cells) and control mice (*n* = 5 mice, 14 cells, KS test, *P* > 0.05). (**G**) Diagram of injection of AAV-hSyn-DIO-ChR2- mCherry plus AAV-VGlut2-Cre-2A- mCherry into VMHvl and representative image of VMHvl neurons expressing *mCherry.* (**H**) Diagram of whole–cell patch-clamp recordings performed in mCherry-labeled VMHvl neuron and representative trace of light-evoked ChR2 currents recorded in a VMHvl neuron. (**I**) Diagram of AAV-hSyn-DIO-GFP plus AAV-VGlut2-mCherry injected into VMHvl of *VGluT2-Cre* mice and representative image of VMHvl glutamate neurons showing overlapping expression of GFP and mCherry. Statistical significance of inter-event intervals and amplitude of mEPSC was determined by Kolmogorov-Smirnov (KS) test. *P* < 0.05 was considered statistically significant with *ns* indicating non-significant, \**P* <0.05, \*\**P* < 0.01, \*\*\**P* <0.001 and \*\*\*\**P* <0.0001.

## REFERENCES

1. J. T. McCracken et al., Risperidone in children with autism and serious behavioral problems. N Engl J Med 347, 314 (Aug 01, 2002).

2. X. Q. Wan et al., Risperidone stimulates food intake and induces body weight gain via the hypothalamic arcuate nucleus 5-HT2c receptor-NPY pathway. CNS Neurosci Ther 26, 558 (May, 2020).

3. H. Lee et al., Scalable control of mounting and attack by Esr1+ neurons in the ventromedial hypothalamus. Nature 509, 627 (May 29, 2014).

4. D. Lin et al., Functional identification of an aggression locus in the mouse hypothalamus. Nature 470, 221 (Feb 10, 2011).

5. C. F. Yang et al., Sexually dimorphic neurons in the ventromedial hypothalamus govern mating in both sexes and aggression in males. Cell 153, 896 (May 9, 2013).

6. D. W. Kim et al., Multimodal Analysis of Cell Types in a Hypothalamic Node Controlling Social Behavior. Cell 179, 713 (Oct 17, 2019).

7. R. Remedios et al., Social behaviour shapes hypothalamic neural ensemble representations of conspecific sex. Nature 550, 388 (Oct 18, 2017).

8. S. C. Rogan, B. L. Roth, Remote control of neuronal signaling. Pharmacol Rev 63, 291 (Jun, 2011).

9. J. T. Glessner et al., Autism genome-wide copy number variation reveals ubiquitin and neuronal genes. Nature 459, 569 (May 28, 2009).

10. M. E. K. Niemi et al., Common genetic variants contribute to risk of rare severe neurodevelopmental disorders. Nature 562, 268 (Oct, 2018).

11. B. K. Bulik-Sullivan et al., LD Score regression distinguishes confounding from polygenicity in genome-wide association studies. Nat Genet 47, 291 (Mar, 2015).

12. D. J. Weiner et al., Polygenic transmission disequilibrium confirms that common and rare variation act additively to create risk for autism spectrum disorders. *Nat Genet*, (May 15, 2017).

13. V. Krishnan et al., Autism gene Ube3a and seizures impair sociability by repressing VTA Cbln1. Nature 543, 507 (Mar 23, 2017).

14. S. E. Smith et al., Increased gene dosage of Ube3a results in autism traits and decreased glutamate synaptic transmission in mice. Sci Transl Med 3, 103ra97 (Oct 5, 2011).

15. R. Anney et al., Individual common variants exert weak effects on the risk for autism spectrum disorders. Hum Mol Genet 21, 4781 (Nov 1, 2012).

16. T. Gaugler et al., Most genetic risk for autism resides with common variation. Nat Genet 46, 881 (Aug, 2014).

17. J. Grove et al., Identification of common genetic risk variants for autism spectrum disorder. Nat Genet 51, 431 (Mar, 2019).

18. L. Klei et al., Common genetic variants, acting additively, are a major source of risk for autism. Mol Autism 3, 9 (Oct 15, 2012).

19. Y. H. Jiang et al., Mutation of the Angelman ubiquitin ligase in mice causes increased cytoplasmic p53 and deficits of contextual learning and long-term potentiation. Neuron 21, 799 (Oct, 1998).

20. T. Matsuura et al., De novo truncating mutations in E6-AP ubiquitin-protein ligase gene (UBE3A) in Angelman syndrome. Nat Genet 15, 74 (Jan, 1997).

21. Z. Nawaz et al., The Angelman syndrome-associated protein, E6-AP, is a coactivator for the nuclear hormone receptor superfamily. Mol Cell Biol 19, 1182 (Feb, 1999).

22. M. Scheffner, J. M. Huibregtse, R. D. Vierstra, P. M. Howley, The HPV-16 E6 and E6-AP complex functions as a ubiquitin-protein ligase in the ubiquitination of p53. Cell 75, 495 (Nov 5, 1993).

23. H. Dhillon et al., Leptin directly activates SF1 neurons in the VMH, and this action by leptin is required for normal body-weight homeostasis. Neuron 49, 191 (Jan 19, 2006).

24. T. Uemura et al., Trans-synaptic interaction of GluRdelta2 and Neurexin through Cbln1 mediates synapse formation in the cerebellum. Cell 141, 1068 (Jun 11, 2010).

25. W. Pereanu et al., AutDB: a platform to decode the genetic architecture of autism. Nucleic Acids Res 46, D1049 (Jan 4, 2018).

26. H. M. Grayton, M. Missler, D. A. Collier, C. Fernandes, Altered social behaviours in neurexin 1alpha knockout mice resemble core symptoms in neurodevelopmental disorders. PLoS One 8, e67114 (2013).

27. R. Yadav et al., Deletion of glutamate delta-1 receptor in mouse leads to aberrant emotional and social behaviors. PLoS One 7, e32969 (2012).

28. L. Lo et al., Connectional architecture of a mouse hypothalamic circuit node controlling social behavior. Proc Natl Acad Sci U S A 116, 7503 (Apr 9, 2019).

29. H. A. Dierick, R. J. Greenspan, Serotonin and neuropeptide F have opposite modulatory effects on fly aggression. Nat Genet 39, 678 (May, 2007).

30. S. L. Padilla et al., Agouti-related peptide neural circuits mediate adaptive behaviors in the starved state. Nat Neurosci 19, 734 (May, 2016).

31. K. Konno et al., Enriched expression of GluD1 in higher brain regions and its involvement in parallel fiber-interneuron synapse formation in the cerebellum. J Neurosci 34, 7412 (May 28, 2014).

32. S. M. Sternson, G. M. Shepherd, J. M. Friedman, Topographic mapping of VMH --> arcuate nucleus microcircuits and their reorganization by fasting. Nat Neurosci 8, 1356 (Oct, 2005).

33. K. Deisseroth, Optogenetics: 10 years of microbial opsins in neuroscience. Nat Neurosci 18, 1213 (Sep, 2015).

34. L. Madisen et al., A toolbox of Cre-dependent optogenetic transgenic mice for light- induced activation and silencing. Nat Neurosci 15, 793 (May, 2012).

35. C. U. Correll, T. Lencz, A. K. Malhotra, Antipsychotic drugs and obesity. Trends in Molecular Medicine 17>, 97 (Feb, 2011).

36. C. A. Doyle, C. J. McDougle, Pharmacologic treatments for the behavioral symptoms associated with autism spectrum disorders across the lifespan. Dialogues Clin Neurosci 14, 263 (Sep, 2012).

37. W. D. Todd et al., A hypothalamic circuit for the circadian control of aggression. Nat Neurosci 21, 717 (May, 2018).

38. L. C. Wong et al., Effective Modulation of Male Aggression through Lateral Septum to Medial Hypothalamus Projection. Curr Biol 26, 593 (Mar 7, 2016).

39. R. M. Quadros et al., Easi-CRISPR: a robust method for one-step generation of mice carrying conditional and insertion alleles using long ssDNA donors and CRISPR ribonucleoproteins. Genome Biol 18, 92 (May 17, 2017).

40. J. M. Koolhaas et al., The resident-intruder paradigm: a standardized test for aggression, violence and social stress. J Vis Exp, e4367 (2013).

41. D. J. Urban, B. L. Roth, DREADDs (designer receptors exclusively activated by designer drugs): chemogenetic tools with therapeutic utility. Annu Rev Pharmacol Toxicol 55, 399 (2015).

42. P. Bankhead et al., QuPath: Open source software for digital pathology image analysis. Sci Rep 7, 16878 (Dec 4, 2017).

43. A. Citri, Z. P. Pang, T. C. Sudhof, M. Wernig, R. C. Malenka, Comprehensive qPCR profiling of gene expression in single neuronal cells. Nat Protoc 7, 118 (Jan, 2012).

